# Subclass IId bacteriocins targeting Man-PTS—structural diversity and implications for receptor interaction and antimicrobial activity

**DOI:** 10.1101/2024.06.22.600115

**Authors:** Aleksandra Tymoszewska, Tamara Aleksandrzak-Piekarczyk

**Author notes:** Corresponding author. Mailing address: IBB PAS, Pawińskiego 5a, 02-106 Warsaw, Poland.

## Abstract

The bacterial mannose phosphotransferase system (Man-PTS) mediates uptake of selected monosaccharides. Simultaneously, it is a receptor for diverse bacteriocins such as subclass IIa pediocin-like bacteriocins and some subclass IId ones (garvicins ABCQ, lactococcins ABZ, BacSJ, ubericin K, and angicin). So far, no attempt has been made to categorize this ever-expanding group of bacteriocins. Here, we identified Man-PTS as a receptor for a number of novel bacteriocins and demonstrated that they all belong to a large family of Man-PTS-binding non-pediocin-like peptides. Based on amino acid sequence similarities between members of this family, we propose their classification into five groups. This classification conveniently distinguishes bacteriocins with specific structures and properties regarding their spectrum of antimicrobial activity and pattern of interaction with Man-PTS. With respect to the latter, we indicate individual amino acid residues or regions of Man-PTS and the bacteriocin responsible for their interaction. In Man-PTS these residues localize to the exterior of the transport complex, specifically the extracellular loop of the so-called Vmotif domain containing regions γ and/or γ+, and to the interior of the transport complex, specifically the interface between the Core and Vmotif domains. Finally, we propose that while the bacteriocins from separate groups display specific binding patterns to Man-PTS, the general mechanism of their interaction with the receptor is universal despite significant differences in their predicted structures, i.e., after initial docking on the bacterial cell through an interaction with the Man-PTS regions γ and/or γ+, they pull away its Core and Vmotif from one another to form a pore across the membrane.

**Significance statement:** Bacteriocins show potential as natural and safe food preservatives and next-generation antibiotics. However, ensuring their safe future use requires primarily the identification of bacteriocin receptors and a detailed understanding of the molecular mechanisms of their selective recognition and binding. Here, we demonstrate the paramount role of Man-PTS in the binding of various non-studied and nearly non-homologous subclass IId bacteriocins with different activity spectra and bacteriocin-receptor binding patterns. Exploiting Man-PTS as a target for novel antimicrobials could be a promising strategy of killing diverse bacterial pathogens, including their antibiotic-resistant strains.

## Introduction

The mannose phosphotransferase system (Man-PTS) is a major glucose and mannose transporting system in *Bacillota* and *Gammaproteobacteria* (1) that can also transport a variety of other hexoses including fructose, glucosamine, *N*-acetylglucosamine, and galactosamine (2). It belongs to phosphoenolpyruvate (PEP)-dependent transport systems which couple the import and phosphorylation of sugars. The transfer of the phosphoryl group from PEP to the incoming sugar is performed sequentially by the cytoplasmic components (enzyme I, HPr, and subunits IIA and IIB of enzyme II) of Man-PTS, while its membrane-localized components (subunits IIC and IID of enzyme II) form a sugar-translocating complex (3). In addition to its function in sugar uptake, Man-PTS regulates numerous cellular processes, e.g., gene expression, metabolism, biofilm formation, and virulence (4). Moreover, the membrane complex of Man-PTS is a receptor for bacteriophage λ (5) and some bacteriocins, i.e., bacterial ribosomally synthesized peptides with bacteriostatic or bactericidal activity (6–12).

Although Man-PTS-targeting bacteriocins are produced by both Gram-negative and Gram-positive bacteria, most of them originate from Gram-positive bacteria and belong to bacteriocins of class II which, in contrast to class I bacteriocins, do not undergo post-translational modifications other than formation of disulfide bridges. Class II bacteriocins are usually divided into four subclasses: IIa pediocins (YGNG-motif containing bacteriocins), IIb two-peptide bacteriocins, IIc leaderless bacteriocins bacteriocins, and IId non-pediocin-like, single-peptide, other linear bacteriocins (non-YGNG-like) (13, 14). Man-PTS is the receptor for pediocin PA-1 (PedPA-1) and most other pediocins (6), as well as subclass IId garvicins Q, A, B, and C (GarQ, A, B, and C), lactococcins A, B, and Z (LcnA, B, and Z), BacSJ, ubericin K, and angicin (6–12). All pediocins share remarkable primary sequence similarity (40-60%) (15), have a conserved amino acid motif (YGNG[V/L]), and a relatively wide activity spectrum (16). In contrast, subclass IId bacteriocins vary considerably in amino acid sequence and activity spectrum, from broad (GarQ, BacSJ, and ubericin K) to narrow (LcnABZ and GarABC) (6–12).

The differences in the bacterial sensitivity to the class II Man-PTS-targeting bacteriocins may result from the presence of unique regions termed α, γ, and γ+ in the Man-PTS membrane complex. Region α localizes to the N-terminal half of the IIC subunit and, depending on the strain, contains a conserved GGQGxxG or GG[D/K]FxxxG motif, while regions γ and γ+ constitute a 35-40 and 51 amino acid stretch, respectively, in the extracellular loop of the IID subunit (8, 16). Pediocins are active against bacteria whose Man-PTS contains regions α and γ, which belong to the genera *Carnobacterium*, *Clostridium*, *Enterococcus*, *Lactobacillus*, *Leuconostoc*, *Listeria*, *Pediococcus*, and *Streptococcus*, and region α determines their sensitivity to pediocins (16). Similarly to pediocins, broad-spectrum IId bacteriocins are active against bacteria with Man-PTS regions α and γ. However, their activity spectrum is even broader and also includes *Lactococcus* spp. GarQ and ubericin K are both active against *Lactococcus lactis* (containing only γ region) and *Lactococcus garvieae* (containing regions γ and γ+), while BacSJ targets *L. lactis* only (7, 10, 11). Among narrow-spectrum IId bacteriocins, LcnABZ are active only against *L. lactis* (6, 9), GarAB against *L. garvieae*, and GarC against all *Lactococcus* spp. (8). In the case of GarABC, region γ+ is required for their activity against *L. garvieae* (8).

Although Man-PTS is recognized as a receptor by several bacteriocins, the exact mechanisms of their action appear to differ. Some of those bacteriocins have been speculated to disrupt the membrane integrity using an unknown mechanism of pore formation (PedPA-1, LcnA, LcnB, ubericin K, angicin, and GarQ) (11, 12, 17–20), while others were postulated to inhibit cell growth (LcnZ) (9) or cell division (GarA) (21), or to have multiple, concentration-dependent mechanisms of action (GarA) (8). Recently, the three-dimensional (3D) structures of Man-PTS components IICIID from *Listeria monocytogenes* and *L. lactis* in complex with PedPA-1 and LcnA, respectively, have been resolved allowing the mechanism of membrane pore formation by these two bacteriocins to be proposed (22, 23). The IICIID complex is a homotrimer with an inward-facing conformation and probably an elevator-type mannose transport mechanism. Each protomer consists of interlocked subunits IIC and IID, which adopt a similar fold of two reentrant loops (hairpins HP1 and HP2), amphipathic helix (AH), and five transmembrane helices (TH1-TH5). The IIC and IID subunits are related to each other by a two-fold pseudo-symmetry axis forming two separate Core and Vmotif domains. The Core domain composed of the HPs, TH1, and TH2 fragments of IICIID is the mannose-binding domain, while the Vmotif domain comprising the TH3-TH5 helices of IICIID forms the scaffold of the trimer. The two domains are connected by AHs. During sugar transport, the Core domain moves vertically across the cell membrane relative to the stationary Vmotif domain (3, 22, 23).

A similar mechanism of membrane pore formation has been postulated for PedPA-1 and LcnA. The both bacteriocins contain three β-strands at the N-terminus and one α-helix at the C-terminus. Initially, the N-terminal domain recognizes the receptor in its outward-facing conformation, attaching to the binding site on the extracellular surface of the Core domain (22, 23). The C-terminal domain then penetrates the IICIID complex and thus the membrane, forming a pore by pulling the Core domain away from the Vmotif domain. However, PedPA-1 and LcnA bind at different positions on the Core domain, which may account for the observed formation of pores with distinct sizes by these two bacteriocins and their strikingly different range of antimicrobial activity (22, 23). It is not yet known whether a similar mechanism is used by other pore-forming bacteriocins targeting Man-PTS.

In this work, we greatly expand the range of bacteriocins recognizing Man-PTS, thereby providing an unequivocal confirmation of the paramount role of this system as the receptor for a variety of non-homologous bacteriocins from subclass IId. According to our comparative analysis, these bacteriocins belong to several distinct and distant groups characterized by specific structural features. These differences likely shape different patterns of interaction with the receptor and thus determine the ranges of bacterial susceptibility. By substituting individual amino acid residues in Man-PTS and selecting those that reduce bacterial sensitivity to particular bacteriocins, we identified distinct binding patterns for each non-homologous bacteriocin, even though they use the same general mode of action. The identification of such a large group of bacteriocins from subclass IId as targeting the same receptor warrants placing them in a new family of Man-PTS-binding non-pediocin-like bacteriocins.

## Results

### Peptides similar to known Man-PTS-targeting bacteriocins form a large family of non-homologous groups

Our earlier studies on GarQ, A, B, C, and BacSJ suggested that Man-PTS could serve as a receptor for a variety of dissimilar subclass IId bacteriocins, which therefore should be classed as a large family of Man-PTS-targeting bacteriocins (7, 8, 10). Here, through a literature review and homology searches, we identified over 40 peptides sharing some amino acid sequence similarity with the aforementioned garvicins and BacSJ (Fig. 1). Among them were bacteriocins known to target Man-PTS (ubericin K, angicin, LcnA, LcnB, and LcnZ) (6, 9, 11, 12), bacteriocins with unknown receptors (bovicin 255 [Bov255], garvicins AG1 and AG2 [GarAG1 and GarAG2]) (24, 25), as well as 33 hypothetical bacteriocins (HB) whose antimicrobial activity and receptors have not been identified. Based on their amino acid sequence similarity, five groups were distinguished and designated GarQ-, LcnA-, LcnB-, GarC- and GarA-like after the first identified representative. Among them, the GarQ- and GarC-like groups were the most numerous with 22 and 12 members, respectively, while the other groups had only 3 or 4 representatives (Fig. 1). The assignment of the bacteriocins to particular groups was mostly confirmed by a phylogenetic analysis (Fig. S1). GarAG1 and HB32 forming an outgroup were a notable exception: despite some convergence of their amino acid sequences, especially at the C-terminus, with the GarA-like peptides, they turned out to be only distantly related to the other Man-PTS-binding bacteriocins. Similarly, HB15 and HB16 were assigned to a phylogenetic group separate from the GarQ-like one and more closely related to the LcnA-like bacteriocins (Fig. S1). The amino acid sequences of HB15 and HB16 show some similarity to the both groups, and their inclusion in the GarQ-like group (Fig. 1) was based on the conserved N-terminus shared with the other bacteriocins in this group.

**Figure 1.**
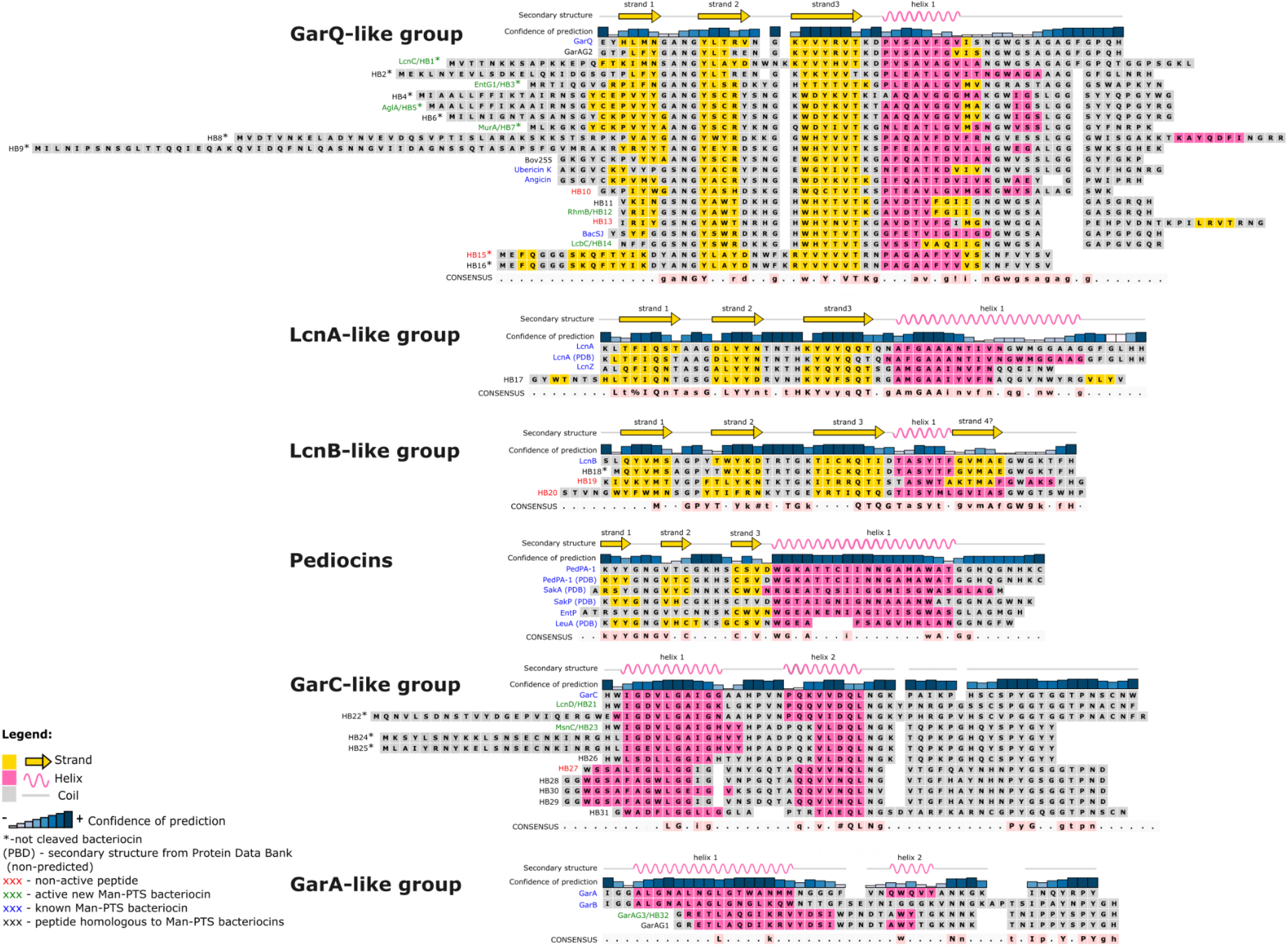
Sequence alignment of peptides showing similarity to Man-PTS-binding bacteriocins. Five novel groups were distinguished based on their amino acid sequences; the pediocin-like group is presented here for comparison. Bacteriocins shown earlier to require Man-PTS for activity are in blue; novel bacteriocins shown here to recognize Man-PTS are in green; hypothetical bacteriocins (HBs) that showed no activity against any indicator strain are in red; HBs that have not been assayed are in black. Consensus symbols are: !—I or V; %—F or Y; #—D, E or N. For the GarQ-like group, the conserved amino acid motifs in the consensus were defined on the basis of biologically active peptides only, for the other groups on the basis of all homologous peptides from the alignment. The amino acid sequences shown are those of mature peptides, after the signal peptide is cleaved off just downstream of the double-glycine motif; if such a motif is absent, a full sequence is shown and marked with an asterisk (*). Secondary structures were predicted *in silico* or derived from a resolved structure deposited in the Protein Data Bank (PDB) repository. The graphical representation of secondary structure elements and the secondary structure prediction confidence index applies only to the first representative of a given group. Strand 1 of Pediocin PA-1 is marked based on the PDB structure and low confidence index of this region.

A secondary structure prediction revealed that all members of a group share a similar pattern within the group but show slightly or significantly different structures between groups. The GarQ-, LcnA- and LcnB-like bacteriocins have a similar general arrangement of predicted secondary structures typical for pediocins, that is, a three-stranded antiparallel β-sheet at the N-terminal part and a single α-helix at the C-terminus. However, compared to pediocins, the three β-strands were much longer and were predicted with a higher confidence. In contrast, their α-helices were typically shorter and of lower confidence than their pediocin counterparts. This is especially evident in the GarQ- and LcnB-like bacteriocins, whose α-helices are substantially shorter and some are even terminated by an additional β-strand. The GarA- and GarC-like bacteriocins have a substantially different regions of predicted secondary structure comprising only two α-helices in the N-terminal part and an unstructured C-terminus. Moreover, their α-helical fragments are not identical; in the GarC-group, both helices are well formed and of confident prediction, while in the GarA-like bacteriocins only the first α-helix is prominent, while the second one is of lesser confidence or apparently not formed at all (Fig. 1).

The above conclusions were supported by a tertiary structure prediction of the main representative of each group. GarQ and LcnB were predicted to have an elongated structure similar to that of pediocins, with two spatially separated parts—a β-sheet at the N-terminal part and an α-helical C-terminal part. In contrast, GarC and GarA were predicted to have a more compact globular structure with one (GarA) or two (GarC) N-terminal α-helices and an unstructured C-terminus (Fig. S2).

### Man-PTS is a receptor for the putative non-homologous bacteriocins

To determine whether Man-PTS is indeed a receptor for the newly identified putative bacteriocins, 15 representatives (HB1, HB3, HB5, HB7, HB10, HB12, HB13, HB14, HB15, HB19, HB20, HB21, HB23, HB27, HB32) (Fig. 1) were synthesized chemically and verified by MALDI-TOF MS. Their antimicrobial activity was then determined against (i) wild-type *L. garvieae* and *L. lactis*; (ii) their mutants with the operons encoding Man-PTS (*manABCD* and *ptnABCD*, respectively) deleted; and (iii) *L. lactis ptnABCD*-deletion strain expressing *manABCD*, *ptnABCD* or their individual genes (Table S2). The activity of nine peptides (HB1, HB3, HB5, HB7, HB12, HB14, HB21, HB23, HB32) depended on the presence of the IICIID complex, thus indicating that they indeed should be classed as Man-PTS-binding bacteriocins. They were therefore given new names based on their host organisms and classified as GarQ-like (lactococcin C [LcnC], enterocin HSIEG1 [EntG1], agilicin A [AglA], murinocin A [MurA], rhamnosin B [RhmB], lactobin C [LcbC]), GarC-like (lactococcin D [LcnD], mesentericin C [MsnC]) and GarA-like (garvicin AG3 [GarAG3]) bacteriocins. The six remaining peptides (HB10, HB13, HB15, HB19, HB20, HB27) proved inactive against any of the *Lactococcus* spp. strains, suggesting that they do not use Man-PTS as a receptor, have no biological activity, or are active against other strains than those tested here (Fig. 1; Table S2).

### The Man-PTS-targeting bacteriocins have a diverse spectrum of antimicrobial activity

The activity of nine newly identified bacteriocins targeting Man-PTS (LcnC, EntG1, AglA, MurA, RhmB, LcbC, LcnD, MsnC, and GarAG3) was tested against a wide range of Gram-positive and Gram-negative bacteria, and yeast. Also ubericin K, angicin, GarQ, GarA, GarB, GarC, and BacSJ were examined in the same manner to expand the spectrum of strains tested earlier (7, 8, 10–12). The bacteriocins assayed turned out to have markedly diverse spectra of activity but limited only to Gram-positive bacteria (Fig. S3).

Bacteriocins from the GarQ-like group showed the broadest spectrum of antimicrobial activity, particularly ubericin K and GarQ which were bactericidal against most strains of the genera *Carnobacterium*, *Enterococcus*, *Lacticaseibacillus*, *Lactiplantibacillus*, *Lactobacillus*, *Lactococcus*, *Leuconostoc*, *Listeria*, *Pediococcus*, and (ubericin K only) *Streptococcus*. Other bacteriocins from this group, such as angicin, BacSJ, MurA, and RhmB had a similarly broad spectrum of antimicrobial activity, except that none was lethal against *L. garvieae* and angicin did not act against *Lactococccus* spp. However, this group also included LcbC, AglA, LcnC, and EntG1 with much narrower spectra of activity limited to strains from the entire genus *Lactococcus* (LcnC), or only *L. lactis* and *L. cremoris* (EntG1), or a few strains of *L. lactis* plus selected strains of species having the α region in their Man-PTS (LcbC and AglA). In addition, a common feature of these four bacteriocins was the absence or negligible activity against *L. monocytogenes*, which was quite unexpected given the strong antilisterial activity of GarQ-like bacteriocins. In the case of AglA, LcnC, and EntG1, this could be due to the presence of an extended N-terminus. These three peptides lacked the protease-processing site with glycines at positions −1 and −2 (relative to the cleavage site) and therefore their signal peptides were not removed. The fourth of these bacteriocins, LcbC, had its signal peptide cleaved off but nevertheless it was also poorly active against *L. monocytogenes*. This deficiency could be due to an inability of the N-terminal fragment to form a stable β-sheet, since LcbC lacks the first β-strand (Fig. 1).

In contrast to the GarQ-like bacteriocins, the other groups had much narrower activities, usually limited to selected *Lactococcus* spp. Of those, GarC and LcnD from the GarC-like group were consistently active against all strains of the genus *Lactococcus*. Unexpectedly, another bacteriocin from this group, MsnC, had a markedly different activity being bactericidal against some strains with Man-PTS with an α region, but showed a weak activity against *L. monocytogenes*, and a selective strain-dependent activity against *Lactococcus* spp. This atypical activity spectrum could be due to the differences in size and predicted secondary structure between MsnC and GarC and LcnD. MsnC is 7 or 9 residues shorter, respectively, and has a more stable first α helix and the second weaker. The remaining bacteriocins had very narrow spectra of activity. Those assayed here from the GarA-like group were only active against *L. garvieae*, while LcnA, LcnB, and LcnZ (from LcnA-like and LcnB-like groups) were active against *L. lactis*, as had been shown earlier (9, 19, 26).

None of the bacteriocins was active against *Apilactobacillus kunkeei*, *Streptococcus mitis*, *Bacillus* spp. or *Staphylococcus* spp. as these bacteria do not encode Man-PTS with regions α, γ and γ+. The bacteriocins assayed also showed no activity against *Pediococcus acidilactici* whose genome contain appropriate Man-PTS-encoding genes, which could be explained by the extended spacer length between the initiation codon and the ribosome binding site in these species negatively affecting the synthesis of the Man-PTS, as proposed before (7).

To determine whether the six peptides (HB10, HB13, HB15, HB19, HB20, HB27), which apparently did not target Man-PTS, had no biological activity whatsoever or were in fact active but not against *Lactococcus* spp., they were additionally tested against a wider panel of indicator strains. Still, no antimicrobial activity was observed. The lack of HB10 activity could be due to the substitution of the highly conserved Tyr22 forming the third β-strand of GarQ-like bacteriocins with Cys, a drastic change of properties. The residue in position 22 could be directly involved in the receptor binding as equivalent residue of the third β-strand of PedPA-1 (which carries Val rather than Tyr) has been shown to interact with Man-PTS from *L. monocytogenes* (22). HB10 also stands out from the other GarQ-like peptides because its C-terminal part is shorter by at least one residue. Also other inactive peptides showed notable differences in length and structure when compared with active bacteriocins from their respective groups—HB13 has an extended C-terminus with an additional β-strand; HB15 lacks the double-glycine site for protease processing and as a consequence the signal peptide is not removed; HB19 and HB20 are markedly different from the active LcnB representative, being 1 or 5 residues longer, respectively, lacking the last β-strand (HB20) or having longer α-helix (HB20) or an additional one (HB19); HB27 the from GarC-like group has a significantly longer distance between its two α-helices (Fig. 1). Notably, circular dichroism (CD) spectroscopy showed that the aforementioned alterations were significant enough to lead to a complete loss of secondary structures in HB15 and HB20, while the other inactive peptides, despite the amino acid sequence changes, were still well-folded (Fig. S4).

### Specific Man-PTS residues condition sensitivity to subclass IId bacteriocins

Earlier studies have indicated that substitutions of certain residues in Man-PTS decrease the sensitivity of the mutants to Man-PTS-dependent bacteriocins such as GarQ, GarA, GarB, GarC, and BacSJ (7, 8, 10), suggesting that they are likely involved in the binding of the bacteriocins. To identify the individual amino acid residues of Man-PTS required for the sensitivity to the newly defined bacteriocins, and thus probably involved in their interaction with the receptor, spontaneous mutants of *L. lactis* IL1403 and *L. garvieae* IBB3403 with resistance to individual bacteriocins and preserved Man-PTS functionality were selected. Out of the nine new bacteriocins, five (GarAG3, RhmB, LcnC, EntG1, and MurA) were potent enough to produce resistant mutants in their presence. In total, 34 resistant *L. lactis* IL1403 mutants were obtained, including three selected in the presence of RhmB, 15 in the presence of EntG1, 2—LcnC, and 14—MurA; 38 resistant *L. garvieae* IBB3403 mutants were obtained in the presence of GarAG3 (Table S1). They were 4- to over 1024-fold less sensitive to the bacteriocin used for the selection than the parental strain and carried, respectively, a combined total of eleven and ten independent mutations in the *ptnCD* genes of *L. lactis* IL1403 and the *manCD* genes of *L. garvieae* IBB3403. Ten mutations in *L. lactis* led to amino acid substitutions in IIC (Gly62Val, Ala110Thr, Met120Ile, and Ala187Val) or IID (Gly126Ser, Ala127Thr, Ser214Leu, Gly221Cys, Tyr223Cys, and Gly229Val), and one led to the deletion of eight amino acids (221-228GlyAlaTyrLeuGluPheProLys) in IID. In *L. garvieae*, eight mutations caused amino acid substitutions in IIC (Gly60Cys, Gly60Val, and Gly188Val) or IID (Arg203Leu, Arg203Cys, Trp256Leu, Gly267Val, and Ser318Leu), and two resulted in deletions of four amino acids in IID (262AsnValValGly265 and 265GlyAsnGlyVal268). Almost all of the obtained mutants were cross-resistant to the other bacteriocins tested but the levels of resistance varied significantly. Similarly, almost all GarQ-, GarA-, GarB-, GarC-, and BacSJ-resistant missense mutants obtained in our previous studies (7, 8, 10) showed varying degrees of cross-resistance to the new Man-PTS-targeting bacteriocins (Tables S3, S4).

To identify additional Man-PTS amino acids involved in the interaction with the new bacteriocins, mutants of *L. lactis* B529 with a preserved Man-PTS functionality were obtained by site-specific mutagenesis with each of the eight Man-PTS*_L. lactis_* amino acids whose counterparts in Man-PTS*_L. monocytogenes_* bind PedPA-1 (22) substituted individually, giving eight *L. lactis* B529 mutants carrying a Ser55Val, Ile59Phe, Val101Phe, Ile105Phe or Tyr198Val substitution in the IIC subunit or Pro115Val, Thr134Val or Trp201Val substitution in IID (Table S1). All these mutants showed a reduced susceptibility to the bacteriocins tested, suggesting that the selected residues are involved in the binding of subclass IId bacteriocins. In the case of the most potent bacteriocins (GarQ, LcnC, and ubericin K), the extent of the sensitivity reduction depended on the amino acid substituted, while a complete loss of susceptibility of the mutants was almost always observed towards bacteriocins with moderate (BacSJ, MurA, angicin, AglA, LcbC, and LcnC) or low (GarC, MsnC, RhmB, and EntG1) activity against *L. lactis* B529 (Table S5).

### The Man-PTS residues involved in the interaction with subclass IId bacteriocins localize to the interface between the Core and Vmotif domains

PedPA-1 and LcnA have been postulated to use a similar mechanism of pore formation involving insertion between the Vmotif and Core domains of Man-PTS (22, 23). Contrarily, Microcin E492 (MccE492), a bacteriocin targeting Man-PTS specifically in Gram-negative bacteria, is believed to attach into a pocket between the Core and Vmotif domains and then to oligomerize and form a pore directly within the membrane (27). Here, we compared the localization of the amino acids in Man-PTSs*_L. lactis/L. garvieae_* required for the sensitivity to subclass IId bacteriocins with the localization of the amino acids involved in mannose uptake or MccE492 binding in Man-PTS*_E. coli_* and of those that bind PedPA-1 in Man-PTS*_L. monocytogenes_*. We observed a considerable overlap in the localization of the Man-PTS amino acids from *L. lactis*, *L. garvieae* and *L. monocytogenes* relevant to the binding of bacteriocins IId and PedPA-1, respectively (Fig. S5), indicating a general similarity in their mechanisms of action. In contrast, the localization of the Man-PTS amino acids relevant to the binding of bacteriocins IId was markedly different from those interacting with mannose or MccE492 in *E. coli* (Fig. S5). In both *L. lactis* and *L. garvieae* most of the amino acids in question were located in the extracellular loop of the IID subunit, in the γ region in *L. lactis* and in the γ and γ+ regions in *L. garvieae*. The remaining residues were found in the transmembrane regions HP2, TH1, and TH3 of subunits IICIID (Fig. S5, Fig. S6).

To determine the spatial location of the Man-PTS residues likely interacting with the bacteriocins, they were mapped onto the tertiary structure of Man-PTS*_L. lactis_* (23). Most of the amino acids localized to the interior of the IICIID transport complex, specifically to the interface between the Core and Vmotif domains created by HP2a of IIC and TH1 and TH3 of IICIID, and the others—to the exterior of the transport complex, specifically the extracellular loop of Vmotif domain containing region γ (Fig. 2A). Some of the relevant amino acids from the extracellular loop of IID*_L. garvieae_* could not be mapped, as Man-PTS from *L. lactis* lacks the γ+ region. When the residues conditioning sensitivity to studied bacteriocins were mapped onto the tertiary structure of Man-PTS*_L. lactis_* solved in complex with LcnA (23), the majority of them localized to the Man-PTS regions surrounding LcnA (Fig. 2B) further suggesting that at least some of the bacteriocins studied here, presumably those structurally similar to LcnA and PedPA-1 (GarQ-like), also use a similar mechanism of pore formation.

**Figure 2.**
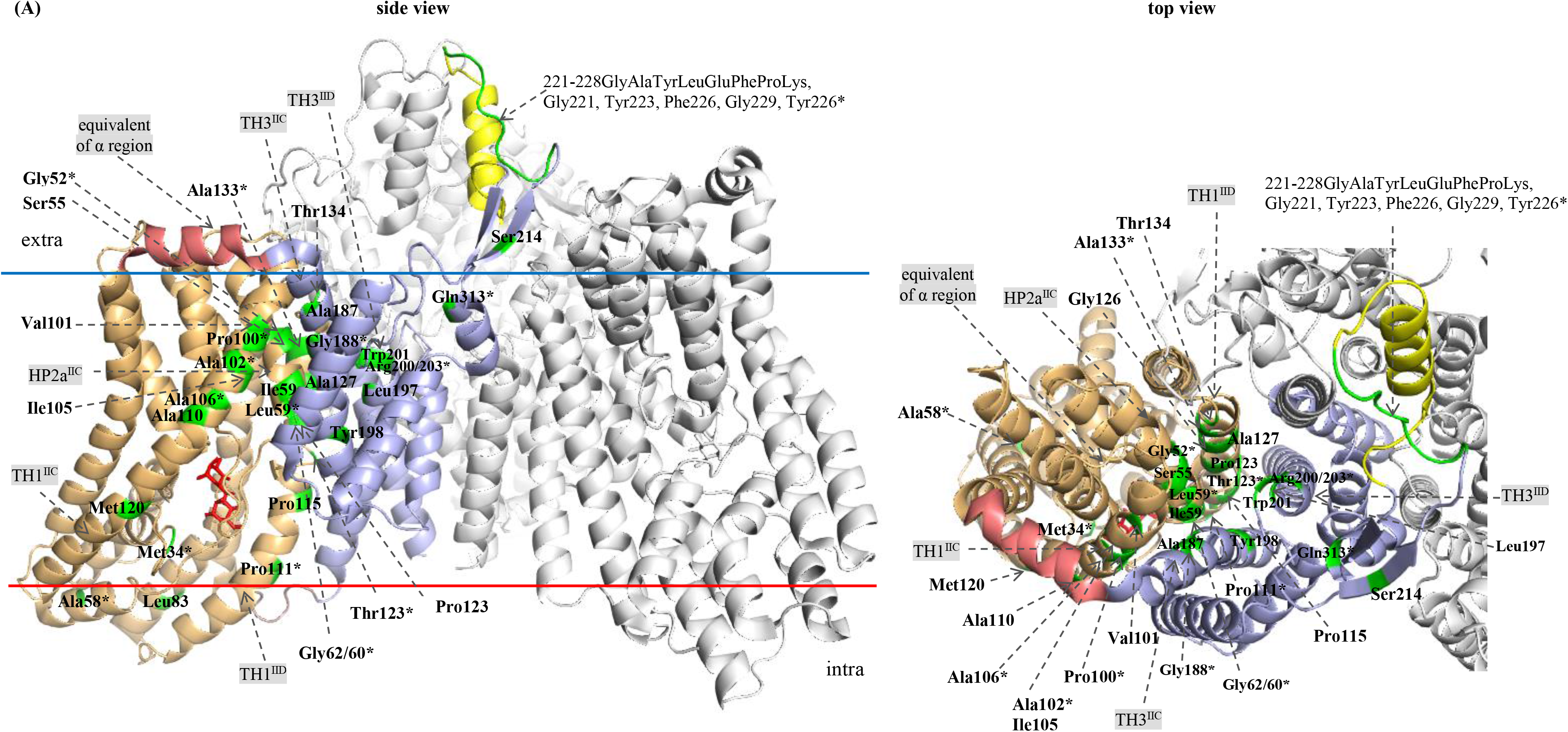
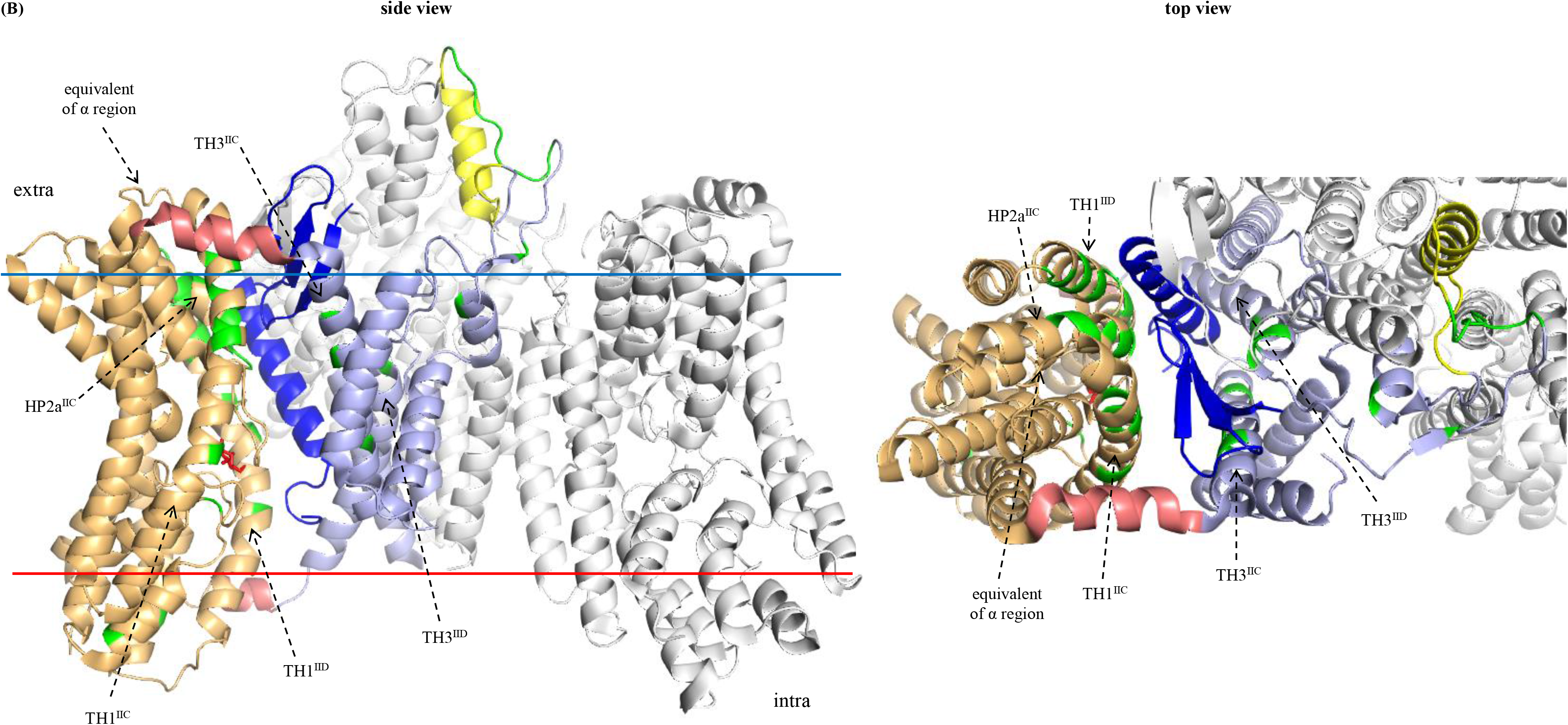
Localization of Man-PTS*L. lactis* amino acid residues required for sensitivity to subclass IId bacteriocins. The amino acids whose spontaneous or site-directed mutations caused resistance to bacteriocins of subclass IId are shown in Man-PTS*L. lactis* (A) and in Man-PTS*L. lactis*-LcnA complex (B). In one Man-PTS protomer the Core domain formed by HPs, TH1 and TH2, the Vmotif domain formed by TH3-TH5, and AH is coloured orange, blue and pink, respectively; region γ in the Vmotif domain is in yellow. Mannose stick model is in red. LcnA is in navy. The Man-PTS amino acids required for sensitivity to bacteriocins are in green. * indicates that the amino acid number refers to Man-PTS*L. garvieae*.

### Man-PTS bacteriocins have diverse but group-specific patterns of interaction with the receptor

Earlier studies have indicated that the extent of sensitivity reduction of a mutant to bacteriocin could reflect a direct (high level) or indirect (low level) involvement of the substituted Man-PTS amino acid in the bacteriocin—receptor interaction (7, 8, 10). Due to the high diversity of the bacteriocins assayed and the differences in their potency, a meaningful comparison of the sensitivity levels of the resistant mutants was challenging. To facilitate the interpretation of the results allowing a robust identification of the Man-PTS amino acids important for bacteriocin binding we used a three-point scale (Fig. 3). Only the spontaneous mutants of *L. lactis* IL1403 and *L. garvieae* IBB3403 were used for the analysis, and bacteriocins with a weak activity against *L. lactis* IL1403 (BacSJ and EntG1) were excluded. Please note that we analyzed here the resistance to all bacteriocins tested, not only those used to select a given mutant.

**Figure 3.**
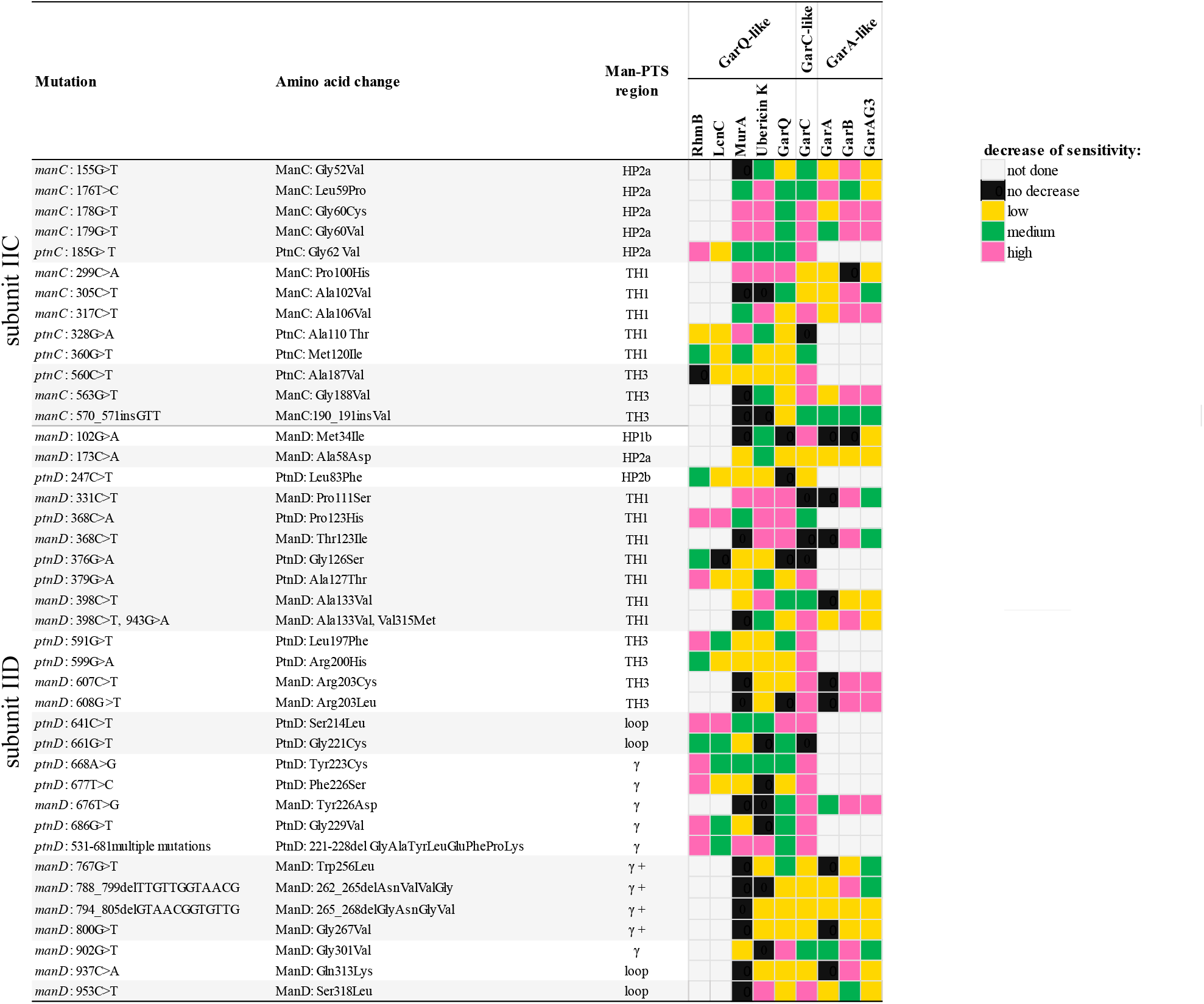
Effect of spontaneous Man-PTS mutations on the sensitivity of *L. lactis* and *L. garvieae* to GarQ-, GarA-, and GarC-like bacteriocins. For each bacteriocin, a fold-decrease of the mutant’s sensitivity relative to WT is shown in a three-point scale: RhmB - low 4x, medium 8x, high >16x; LcnC - low 2-8x, medium 16-32x, high >128x; MurA - low 4-32x, medium 64-256x, high >1024x (*L. lactis*) or low 2x, medium 4-8x, high >16x (*L. garvieae*); GarQ - low 2-8x, medium 16-128x, high ≥256x; ubericin K - low 2-8x, medium 16-64x, high >128x (*L. lactis*) or low 2-4x, medium 8-16x, high >32x (*L. garvieae*); GarA - low 2-4x, medium 8x, high 32x; GarB - low 4-8x, medium 32x, high ≥64x; GarC - low 4x, medium 8x, high >32x (*L. lactis*) or low 2-8x, medium 16-64x, high >128x (*L. garvieae*); GarAG3 - low 2-32x, medium 64-256x, high ≥1024x (details in Tables S3 and S4). “Not done” applies to RmnB and LcnC inactive against *L. garvieae*, and to GarA, GarB, and GarAG3 inactive against *L. lactis*.

We observed that the transmembrane regions HP2a of IIC, and TH1 of IIC and IID are similarly important for the interactions with all the bacteriocins in both *L. lactis* and *L. garvieae*, but within these regions specific amino acids appear to be more relevant for certain groups of bacteriocins. For instance, Pro100 of ManC is mainly used for binding by GarQ-like bacteriocins (MurA, ubericin K, and GarQ) and Ala106 of ManC—by GarC and GarA-like bacteriocins (GarB and GarAG3) (Fig. 3). This could be expected, since the Pro100His substitution was selected in the presence of GarQ, and Ala106Val—in the presence of GarC. The two affected positions lie in the same TH1 IIC region (Fig. 2A), suggesting some specificity in the binding pattern of bacteriocins from different groups to particular amino acid residues. On the other hand, TH3 of IIC and IID turned out to be of little relevance to GarQ-like bacteriocins (RhmB, LcnC, MurA, ubericin K, and GarQ) but important for GarC and the GarA-like ones (GarB and GarAG3) (Fig. 3). Among the latter group, GarA did not show a similar dependence, in accordance with its behavior in several other cases as well, probably caused by its dual mode of action (8, 21). In particular, Gly188 of ManC and Arg203 of ManD (substitutions of which were selected in the presence of GarAG3) were important for the binding of GarC and GarA-like bacteriocins. These two amino acids lie inside the Man-PTS membrane complex, between its Vmotif and Core domains (Fig. 2A), further suggesting that the broad (GarQ-like) and narrow (GarA-like, GarC) spectrum bacteriocins use different binding patterns even though the two types both insert between these two Man-PTS domains.

Likewise, we observed a high significance of the transmembrane HP1b region of IID for the sensitivity to GarC and no such significance for the GarQ- and GarA-like bacteriocins (Fig. 3), further confirming that GarC does differ in its binding pattern from other Man-PTS-binding bacteriocins. The Met34Ile substitution in the HP1b region obtained under GarC selection unexpectedly lies in the mannose-binding pocket (Fig. 2A), suggesting that its role in conferring resistance could be due to some as yet undefined mechanism rather than to its direct effect on the bacteriocin binding.

As to the relevance of the extracellular Man-PTS region for the bacteriocin sensitivity of *L. lactis* and *L. garvieae*, the resistance levels of mutants with altered amino acids in the extracellular loop of the IID subunit indicated that the γ region is important for the interaction with bacteriocins from all groups, especially its N-terminal and middle parts, which were much more frequent hotspots for changes than the C-terminal fragment (Fig. 3). Tyr223 and the eight amino acids at positions 221-228 of PtnD were the most important for all bacteriocin groups, while Tyr226 of ManD was slightly more important for the GarA- and GarC-like groups. All these amino acids are in close proximity to the interface of the Vmotif and Core domains, indicating that they could serve as an initial binding site allowing the bacteriocin to dock on the receptor before inserting between the two Man-PTS domains (Fig. 2A). On the other hand, region γ+ is almost irrelevant for the GarQ-like bacteriocins, which is to be expected given their broad spectrum of activity which also includes bacteria lacking the γ+ region, but appears important for GarB and GarAG3, which both target only *L. garvieae* (Fig. 3). Within the γ+ region, four amino acids at positions 262-265 of ManD were especially important for these two GarA-like bacteriocins. This suggests that the GarA-like bacteriocins share a similar receptor binding pattern, significantly different from that of other bacteriocin groups, in which region γ+ is the main determinant of strain sensitivity, being responsible for the initial receptor recognition.

### The GarQ-like bacteriocins carry conserved amino acid motifs

A comparison of the amino acid sequences of bacteriocins with a proven biological activity from the most abundant GarQ-like group revealed the presence of conserved motifs, such as gaNGY, YxVTK, and nGw, in their N-terminal, central, and C-terminal parts, respectively. Prediction of the secondary and tertiary structures of GarQ showed that the gaNGY and YxVTK motifs are located in the β-sheet of the N-terminal part, while the nGw motif is lies the C-terminal part right at the end of the α-helix (Fig. 1; Fig. S2). The N-terminal part of PedPA-1 containing the YGNG[V/L] motif has been shown to recognize the Man-PTS receptor in a broad spectrum of bacteria containing region α (22), suggesting that the gaNGY and YxVTK motifs of GarQ may have the same function, especially the first one, which is located similarly to the pediocin motif, i.e. in the loop linking the first two β-strands (Fig. 1).

A comparison of two subgroups of GarQ-like bacteriocins: (i) GarQ, GarAG2, LcnC, MurA, and ubericin K, which are all active against two lactococcal species *L. lactis* and *L. garvieae*, with (ii) RhmB, BacSJ, LcbC, angicin, AglA, and EntG1, bactericidal only against *L. lactis*, revealed significant C-terminal conservation among the representatives of the former group and its absence in the latter. The motif that distinguishes these two subgroups lies in the C-terminal part of the α-helix and in the unstructured C-terminus and has the glycine-rich consensus of AvxgVisNGWxgSaGaG (Fig. S7). The presence of this conserved motif in a subgroup of GarQ-like bacteriocins active against *L. garvieae* suggests that their C-terminal fragment could take part in the recognition and initial interaction with the *L. garvieae* Man-PTS.

### The conserved GarQ amino acid residues are important for its antibacterial activity

GarQ was selected as a representative of the GarQ-like bacteriocins to investigate the role of their conserved motifs by substituting individual residues. To verify that the resulting variants are properly folded we used CD spectroscopy. As expected from their short linear amino acid sequence and earlier structural studies on pediocins (28), which were random coil in an aqueous milieu, also in the case of GarQ the membrane-mimicking trifluoroethanol (TFE) solvent had to be added to induce its helicity. The greatest enhancement in the helicity was observed in 50% TFE (Fig. S8A), and this concentration was used for further studies. In those conditions, all GarQ variants synthesized were still correctly folded (Fig. S8B). The activity of the GarQ variants was tested against a wide range of Gram-positive and Gram-negative bacteria, and yeast, and compared with that of GarQ. Most variants - GarQ_Thr24Ala_, GarQ_Lys25Ala_, GarQ_Asn37Gly_, and GarQ_Gly38Ala_ - showed unchanged spectra of activity, albeit the extent of inhibition of bacterial growth was slightly lower than that shown by native GarQ. However, for some variants the activity spectra were altered markedly. A complete loss of activity against all indicator strains was observed for GarQ_Gly10Ala_, and GarQ_Asn9Gly_ and GarQ_Tyr11Leu_ showed a complete or partial loss of activity towards some strains only (Fig. S3). To quantify the effects of the substitutions on the GarQ activity, minimum inhibitory concentrations of the variants were determined against *L. garvieae*. Substitutions of the highly conserved NGY residues from the gaNGY motif had a dramatic effect on the activity of GarQ, leading to a complete (Gly10) or nearly complete (Asn9 or Tyr11; >512-fold decrease) loss of activity, while a replacement of residues from the YxVTK or nGw motifs only caused, respectively, a moderate (8-32-fold) or weak (2-8-fold) decrease in activity (Fig. 4).

**Figure 4.**
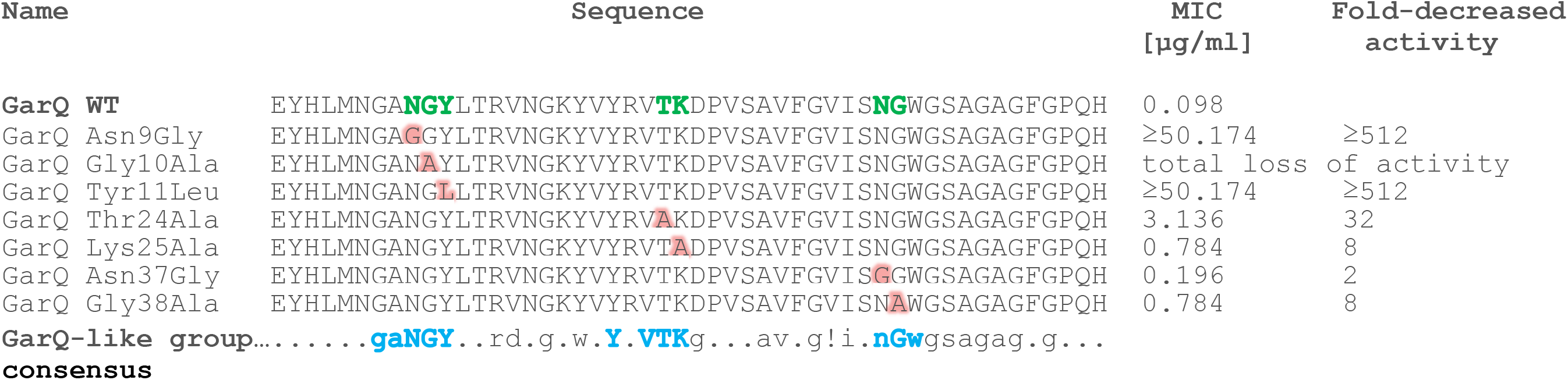
Minimum inhibitory concentrations of GarQ variants against *L. garvieae* IBB3403. In each variant one residue, shown in red, was substituted. All these positions are marked in native (WT) garvicin in green. The most conserved motifs in the entire GarQ-like group are shown in the consensus sequence in blue. The consensus was determined for biologically active GarQ-like peptides only. ! signifies I or V.

## Discussion

The use of bacteriocins as antibacterial agents offers a promising solution to some of the most challenging public health problems today, such as the emergence of difficult-to-treat infections caused by antibiotic-resistant pathogens (29) or the breakouts of foodborne diseases (30). Here, we have identified nine new bacteriocins of which five (MurA, MsnC, RhmB, AglA, and LcbC) could have potential in the treatment of infections caused by vancomycin-resistant *Enterococcus faecium* and *E. faecalis* strains (31), and four (MurA, MsnC, RhmB, and AglA) in the prevention of *L. monocytogenes*-associated food spoilage or treatment of foodborne listeriosis (30). Additionally, some of the newly identified bacteriocins show activity against *Lactococcus garvieae* (MurA, MsnC, LcnC, LcnD, and GarAG3) and *Streptococcus parauberis* (MurA), which indicates their potential in the treatment of animal mastitis, lactococcosis, and streptococcosis (32–35). We showed experimentally that the activity of all bacteriocins studied requires the presence of the Man-PTS membrane complex which most probably serves as their receptor. Man-PTS is absent in eukaryotic cells and highly purified GarQ (at concentrations up to 1 mg/ml) and PedPA-1 (up to 400 μg/ml, a concentration as much as 4,000-fold higher than its MIC for *Listeria innocua*) have already been shown to have no cytotoxicity against Vero (African green monkey kidney), HepG2 (human liver cancer) and Caco-2 (human colon adenocarcinoma) cell lines (36, 37).

It has previously been shown that Man-PTS is the receptor for some bacteriocins of subclasses IIa and IId (6–12). However, those studies were usually limited to a single bacteriocin or a small group of homologous bacteriocins and did not provide information on the range and diversity of bacteriocins interacting with Man-PTS. Based on their amino acid sequence similarity, we distinguished five groups within subclass IId of Man-PTS-binding bacteriocins, namely GarQ-, LcnA-, LcnB-, GarC-, and GarA-like ones, and showed that members of each group are similar to each other at the level of primary and predicted secondary structures, but differ significantly from members of other groups. These differences corresponded with the types of Man-PTS systems targeted by each group. The GarQ-like bacteriocins having a predicted secondary structure similar to that of pediocins (N-terminal part composed of two or three short β-strands and C-terminal part with one α-helix) also had a pediocin-like activity against bacteria containing region α in their Man-PTS. However, their β-strands composed of 3-7 amino acids each were significantly longer than the short β-strands of pediocins (2-4 amino acids long), as well as their entire β-sheet region is more sizeable (Fig. S9), which may explain why the GarQ-like bacteriocins, unlike pediocins, were also active against *Lactococcus* spp. One of the main differences between the listerial and lactococcal Man-PTS is the external surface of their Core domain. In *L. monocytogenes* this surface contains region α localized between HP2b and TH1 of IIC and spatially close to the pediocin-binding residues. In contrast, Man-PTS from *Lactococcus* spp. lacks region α and the corresponding region on the surface of Core domain is two residues longer, preventing the specific binding of the N-terminal PedPA-1 fragment to its recognition site (23). It seems likely that the longer N-terminal β-strands present in the GarQ-like bacteriocins better fit the larger surface of the Core domain of Man-PTS*_L. lactis_* and allow these bacteriocins to reach their binding site.

In contrast to the GarQ-like and PedPA-1-like groups, bacteriocins with long N-terminal β-strands composed of 5-7 amino acids and a C-terminal α-helix (LcnA- and LcnB-like) and those of only an α-helical structure (GarC- and GarA-like) (Fig. S9) were active only against bacteria with Man-PTS lacking region α. This could be due to the long β-strands or α-helices not fitting the external surface of the Core domain of Man-PTS*_L. monocytogenes_*, which contains region α. The only exception to this rule was MsnC from the GarC-like group, active against some strains with Man-PTS with an α region and exhibiting selective strain-dependent activity against *Lactococcus* spp. MsnC contains a strongly developed first α-helix and a weaker second one in comparison with other GarC-like bacteriocins, which may lead to a poorer structural match with Man-PTSs without the α region and a better fit to those with such a region. These results suggest that the N-terminal part of the Man-PTS-targeting bacteriocins is the main determinant of their activity against bacteria containing region α. This would explain why GarQ-like bacteriocins with the N-terminus blocked by the retained signal peptide (AglA, EntG1 and LcnC) or without a properly formed β-sheet (LcbC) show reduced activity against bacteria containing region α but are nevertheless active against *Lactococcus* spp. A similar observation confirming the importance of the free N-terminus of a bacteriocin for binding to the α-region was recently documented by van Belkum et al. who showed that anti-listerial PedPA-1, after blocking its N-terminus by fusion with maltose-binding protein (MBP), lost activity against *L. monocytogenes*, while GarQ and LcnA being in the same fusion were still lethal against *L. lactis* (38). Research at this stage is still limited due to the lack of solved tertiary structures of Man-PTS-binding bacteriocins. Of this group, only two pediocins SakP and LeuA have deposited 3D structures in the PDB repository (entry IDs: 1OG7 and 3LEU, respectively), while the structures of LcnA, PedPA-1 and SakA have been resolved in complex with Man-PTS (entry IDs: 8HFS, 7VLY and 7XNO, respectively) and additionally in N-terminal fusion with large MBP (22, 23, 39), which raises a question about the accuracy of their models (38). Consequently, due to the lack of appropriate templates in PDB, the predictions of the tertiary structures of the tested bacteriocins may have low confidence of prediction. Until 3D models are resolved, research must rely on primary sequences and reliable predictions of secondary structures of bacteriocins, which was achieved in this work.

To study further the interactions between Man-PTS and subclass IId bacteriocins we identified the Man-PTS amino acid residues or regions responsible for its interaction with bacteriocins from the GarQ-, GarA-, and GarC-like groups. We used spontaneous or site-directed mutagenesis of the Man-PTS system which preserved its functionality and allowed identification of a number of residues whose substitution or deletion caused a resistance to the bacteriocins studied, indicating their likely role in the bacteriocin binding. Despite the fairly high number of mutated sites, several mutational hot-spots were apparent, such as the extracellular γ and γ+ regions in the IID subunit, and the transmembrane HP2, TH1, and TH3 regions in the IICIID subunits. The hot-spots in the extracellular regions localized to the exterior loop of the transport complex, specifically its Vmotif domain. In contrast, the hot-spots in the transmembrane regions of IICIID subunits localized to the interior of the Man-PTS membrane complex, specifically the interface between the Core and Vmotif domains. A similar localization of residues involved in PedPA-1 and LcnA binding was previously identified in, respectively, *L. monocytogenes* and *L. lactis* (22, 23), strongly suggesting that all bacteriocins interacting with Man-PTS, both from subclass IId and IIa, use a common mechanism of action.

It is tempting to speculate that the bacteriocins studied initially recognize the sensitive cells by specifically binding to the extracellular loop of the Vmotif domain, predominantly within or adjacent to the γ and γ+ regions. Then, similarly to PedPA-1 and LcnA (22, 23), these bacteriocins recognize selected residues of HP2, TH1 and TH3 localized adjacent to the external surface of the Core domain which in *L. monocytogenes* contains region α while in *Lactococcus* spp. is two residues longer and lacks α region. Finally, the bacteriocins insert between the Vmotif and Core domains by binding to regions of TH1 and TH3 on the interface of these two domains. While inserting, the bacteriocins move the Core domain away from the Vmotif domain forming a pore through the membrane, which finally leads to the efflux of intracellular solutes and consequently cell death. This mechanism differs greatly from the one used by MccE492 which merely uses Man-PTS to anchor on the membrane and forms a pore directly within it after self-oligomerization (27). Not surprisingly, we did not observe any similarity between the localization of the Man-PTS residues involved in the binding of MccE492 and subclass IId bacteriocins.

Although PedPA-1 and LcnA share a common mechanism of pore formation, they recognize and bind at different positions on the Core domain, and consequently adopt a different orientation in the membrane (22, 23). Here, we have identified Man-PTS regions and/or individual amino acids directly involved in the binding of each bacteriocin group by finding those whose substitution or deletion resulted in a moderate or high decrease of the mutant’s sensitivity to the bacteriocin(s) in question. We observed that some Man-PTS regions such as HP2a and γ region are equally important for the sensitivity to all bacteriocins while others, such as HP1b, TH1, TH3 and γ+ region, are of variable importance to individual bacteriocin groups. Thus, TH1 of IICIID, especially Pro100 of ManC and Pro123 of PtnD, are the most important for the interaction with GarQ-like bacteriocins, while TH1 and TH3 of IICIID, especially Ala106 and Gly188 of ManC, and Arg203 of ManD,—with GarC-like bacteriocins. Finally, the same regions of TH1 and TH3 as in the case of GarC-like bacteriocins, and additionally HP1b (especially Met34 of ManD) and γ+ region are the most important for the receptor interaction with GarA-like bacteriocins. Met34 is a unique residue among those mentioned above as it localizes to the mannose binding pocket and is therefore unlikely to be directly engaged in an interaction with the much larger bacteriocins. A similar localization of several other amino acid residues altered by mutations in both the IIC and IID subunits forming the sugar-binding pocket that led to the acquisition of resistance to LcnA and GarQ was presented recently by van Belkum et al. (38). We speculate that these subsites in the sugar-binding pocket through structural changes affect the interface between the Vmotif and Core domains, hindering or even preventing the insertion of bacteriocins. Substitutions of the other amino acids located near the mannose binding pocket (Ala58, Leu83, Met120) did not have such a drastic effect on the mutant sensitivity, suggesting their weak influence on the Man-PTS structure. Based on these results, it is apparent that each bacteriocin group uses a specific Man-PTS binding pattern, slightly or significantly different from those used by other groups, and adopts a different orientation in the membrane.

Also within the group of GarQ-like bacteriocins we found two clear-cut subgroups using different Man-PTS regions for docking: (i) GarQ, MurA, ubericin K, and LcnC, active against *L. lactis* and *L. garvieae*, and (ii) RhmB, BacSJ, LcbC, EntG1, angicin, and AglA, active against *L. lactis* only. The main difference between the Man-PTSs of these two species is the presence of the γ+ region in *L. garvieae* and its absence in *L. lactis*. Therefore, it is tempting to speculate that the initial interaction with γ+ is the main determinant of the bacteriocin activity against *L. garvieae*. On the other hand, mutations in the γ+ region caused almost no changes in the sensitivity to the GarQ-like peptides from the first subgroup, suggesting that in fact they do not bind directly to the γ+ region but rather to the γ region or the extracellular loop (specifically, Ser214, Tyr223, 221-228GlyAlaTyrLeuGluPheProLys, Gly301) which, depending on the bacterial strain, may be accessible or shielded by the presence of the γ+ region. Based on these observations we propose two patterns of docking on the receptor. GarQ, MurA, ubericin K, and LcnC would dock via weak interactions with γ+ and stronger ones with the γ region. Interestingly, MurA, ubericin K, and GarQ exhibit, respectively, weak, moderate, and high potency against *L. garvieae*. These differences correspond with the importance of the γ+ region for their activity, which was the lowest for MurA and the highest for GarQ, with Trp256 being the only residue involved in GarQ binding. The other GarQ-like bacteriocins, included here in subgroup II, RhmB, BacSJ, LcbC, EntG1, angicin, and AglA, are expected to dock using the γ region only, which would explain their lack of activity against *L. garvieae* in which region γ+ constitutes a steric hindrance.

We posit that the conserved amino acid motifs shared by GarQ-like bacteriocins are of key importance for the bacteriocin—receptor interaction. This hypothesis is based on the observation that they are localized in the easily accessible/protruding parts of the bacteriocins and their substitutions decrease the bacteriocin activity without markedly affecting their 3D structure. The largest decrease in activity was observed when residues of the N-terminal NGY motif of GarQ were substituted, suggesting that they are crucial for the specific interaction with the receptor. It is therefore tempting to propose that the NGY motif of the GarQ-like peptides is involved in the binding and stabilization of bacteriocins on the Core domain which, depending on the strain, contains or lacks the α region. This hypothesis is further supported by the similar localization of the NGY motif in GarQ and the Man-PTS-binding YGNG[V/L] motif in PedPA-1. On the other hand, substitutions in the C-terminal nGw motif of GarQ had only a small effect on the bacteriocin activity, suggesting its minor input into the receptor binding. We propose that the C-terminal part of the GarQ-like peptides, which shares high similarity with the transmembrane region of sodium:proline solute transporter (7), could have two functions—it enables the initial docking on the extracellular loop of the Vmotif domain and then, owing to its low binding strength, detaches from the extracellular loop and wedges between the Vmotif and Core domains forming a pore in the membrane. Importantly, the nGw motif may be sufficient for a bacteriocin to dock on to the γ region of *L. lactis*, while the longer AvxgVisNGWxgSaGaG motif may be required for the interaction with the γ+ and γ regions in *L. garvieae*.

Our studies show that the GarQ-like bacteriocins are the most diverse group of bacteriocins targeting Man-PTS and functionally can be divided into at least two markedly different subgroups. Following the present identification of new Man-PTS-targeting bacteriocins it seems almost certain that novel representatives will soon be added to the proposed groups or new groups may have to be formed, such as the GarAG3-like peptides distinguished by the phylogenetic analysis. Although Bov255, GarAG1, and GarAG2 were not studied experimentally in the present study, they seem highly likely to target Man-PTS as well as they have the appropriate structure. The broad activity spectra of Bov255 comprising *Enterococcus* and *Ligilactobacillus* spp., and of GarAG2 comprising *Carnobacterium*, *Enterococcus*, *Lactococcus*, *Listeria*, and *Streptococcus* spp. justify their classification in the GarQ-like group, while the GarAG1 activity limited to *L. garvieae* strains warrants its placement in the GarA-like group of Man-PTS-targeting bacteriocins (24, 25).

In conclusion, by pointing out the paramount role of Man-PTS as the receptor for many homologous and non-homologous bacteriocins with different activity spectra, we offer here a new perspective on the diversity of the cellular receptors for bacteriocins. Despite the large number and variety of subclass IId bacteriocins, Man-PTS is their only identified receptor, with the single exception of lactococcin 972 targeting lipid II (40), strongly suggesting that the diversity of bacteriocin receptors could in fact be much more limited than once postulated. We posit that such an important role of Man-PTS reflects its wide distribution among bacteria from different genera and at the same time a species-specific structure that allowed a co-evolution of a plethora of bacteriocins with different receptor binding patterns and a conserved overall mechanism of pore-formation. A potential reason that Man-PTS is a multitarget for non-homologous bacteriocins may be its specific two-subunit and two-domain structure, unique among other membrane proteins, and which allows the formation of a lethal pore upon contact with a bacteriocin. Based on the lack of homology between the bacteriocin groups, it’s tempting to also speculate that they evolved independently to target this receptor. The presented results markedly expand the understanding of the biology of bacteriocins and foretell novel strategies for future studies and, importantly, possible therapeutic approaches. Homology search has shown its great potential for the identification of multiple previously unknown bacteriocins interacting with a single receptor, while the determination of the bacteriocin-receptor binding patterns can lead to structure-based strategies for designing new antimicrobial peptides with an increased potency or activity against selected bacterial species.

## Materials and Methods

### Bacterial and yeast strains and culture conditions

The microorganisms used in this study are listed in Table S1. Indicator strains were grown in Brain Heart Infusion (BHI) medium (Oxoid, UK) except for *Apilactobacillus*, *Lacticaseibacillus*, *Lactiplantibacillus*, *Lactobacillus*, and *Ligilactobacillus* strains, which were grown in MRS Broth (Oxoid), and *Campylobacter* strains, which were grown on Blood Agar Base No. 2 (Oxoid). *L. garvieae* IBB3403- and *L. lactis* IL1403-derived strains were grown in BHI medium. *Escherichia coli* EC1000-derived strains were grown in Luria-Bertani (LB) medium (Becton, Dickinson and Company, USA).

*Apilactobacillus*, *Lacticaseibacillus*, *Lactiplantibacillus*, *Lactobacillus*, and *Ligilactobacillus* spp. were cultured under anaerobic conditions without shaking at 37°C. *Enterococcus, Pediococcus*, *Streptococcus* spp., and *Candida albicans* were cultured under aerobic conditions without shaking at 37°C. *Lactococcus*, *Leuconostoc*, and *Weissella* spp. were cultured under aerobic conditions without shaking at 30°C. *L. monocytogenes*, *Pseudomonas aeruginosa*, *Salmonella typhimurium* and *Bacillus*, *Escherichia*, and *Staphylococcus* spp. were cultured under aerobic conditions with shaking at 37°C. *Campylobacter* spp. were cultured under microaerobic conditions without shaking at 37°C. *Carnobacterium maltaromaticum* was cultured under aerobic conditions without shaking at 16°C.

To grow *E*. *coli* EC1000-derived strains with pNZ8037, chloramphenicol was added to 20 µg/ml and for *L*. *lactis* IL1403-derived strains with pNZ9530 and pNZ8037, erythromycin and chloramphenicol were added to the growth medium to 5 µg/ml each. Expression of the genes cloned under the nisin-responsive promoter P*_nisA_* in pNZ8037 was induced by the addition of nisin to 10-50 ng/ml. Soft agar or agar plates were prepared by adding agar (Merck, Germany) to 0.75% or 1.5% to the liquid medium, respectively.

### Bacteriocin preparation and MS analysis

Lyophilized peptides with a purity of over 90% were synthesized chemically (PepMic, P.R. China). HB10, RhmB/HB12, HB13, LcbC/HB14, HB19, HB20, LcnD/HB21, MsnC/HB23, HB27, and GarAG3/HB32 were synthesized in mature form. LcnC/HB1, EntG1/HB3, AglA/HB5, MurA/HB7, and HB15, lacking the double-glycine (GG) motif, were synthesized in full form. Leader peptides were predicted using searches of only the basic GG motif, alternative cleveage motifs such as GS, GA, SG or AG were not considered. Stock solutions (1 mg/ml) were prepared by dissolving the peptides in 0.1% trifluoroacetic acid (TFA) (Sigma, Germany). The mass correctness and absence of degradation of the synthetic peptides were verified by matrix-assisted laser desorption ionization time-of-flight mass spectrometry (MALDI-TOF MS) using a two-layer method (41) with 3,5-dimethoxy-4-hydroxycinnamic acid (sinapinic acid; SA) as the matrix. Spectra were acquired in positive ion mode using a Perspective Biosystems Voyager Elite MALDI-TOF MS spectrometer (AB Sciex, USA) in reflectron mode with delayed extraction.

### Circular dichroism spectroscopy

CD spectra were recorded on an OLIS DSM 17CD spectropolarimeter (Bogart, USA) at 20°C in a quartz cell with a 0.02-cm path length over 190-250 nm. The peptide concentration was 0.5 mg/ml. Initially the CD spectra were recorded in 100% H_2_O, 75% H_2_O and 25% TFE, and 50% H_2_O and 50% TFE, and the results obtained were used to optimize the conditions; the latter solvent gave the most pronounced ellipticity signal and was then used for all measurements. Data were recorded every one nm and the results of five scans for each sample were averaged. The bandwidth was set at 2.0 nm. Prior to calculating molar ellipticities, the baseline CD spectrum of the solvent was subtracted from the sample spectra. A digital filter of 11 was applied to the CD spectra. The α-helical content of the peptide was calculated from θ_222nm_ using the equation: % α-helix = (−[θ_222nm_] + 3000)/39000 × 100% (42).

### Detection of bacteriocin activity and selection of bacteriocin-resistant mutants

The activity of peptides of interest against indicator strains and *L. garvieae* IBB3403- or *L. lactis* IL1403-derived strains was determined using the spot-on-lawn assay as described previously (7). Spontaneous bacteriocin-resistant mutants were selected by growing *L. garvieae* IBB3403 or *L. lactis* IL1403 in the presence of a bacteriocin in Chemically Defined Medium (CDM) (43) supplemented with 1% mannose (Man-CDM) as described previously (7). GarAG3-resistant *L. garvieae* IBB3403 mutants (LG1— LG32, LG34—LG38, LG40) were selected at the bacteriocin concentration of 0.024 µg/ml. *L. lactis* IL1403 mutants resistant to EntG1 (LLG1—LLG3, LLG7—LLG9, LLG11, LLG12, LLG14, LLG19, LLG21, LLG25, LLG32, LLG46, LLG50), RhmB (LLB2, LLB3, LLB17), MurA (LLA1, LLA7, LLA8, LLA12, LLA19—LLA22, LLA24, LLA26—LLA30), and LcnC (LLC1, LLC2) were selected at bacteriocin concentrations of 12.5 µg/ml, 1.56 µg/ml, 1.56 µg/ml, and 4 ng/ml, respectively. The sensitivity of the mutants was determined using two-fold serial dilutions assay as described previously (7).

### Sequencing of bacteriocin-resistant mutants

Genomic DNA of *L. garvieae* IBB3403, *L. lactis* IL1403 and their bacteriocin-resistant mutants was extracted using Genomic Mini Kit (A&A Biotechnology, Poland). Samples for sequencing of the *manCD* or *ptnCD* genes were prepared by PCR with *manC*for/rev and *manD*for/rev or *ptnC*for/rev and *ptnD*for/rev primer pairs (Table S1) using the appropriate genomic DNA as a template. PCR was carried out as described previously (10). PCR products were purified using Wizard^®^ SV Gel and PCR Clean-Up System (Promega) and sequenced using Sanger method (44).

### Bioinformatic analysis

The sequencing results were analyzed using Clone Manager software (Sci-Ed, USA). The nucleotide sequences of the *manCD* and *ptnCD* genes were translated to amino acid sequences using the Translate tool on the ExPasy online server (45). The amino acid sequences of ManCD (IICIID*_L. garvieae_*), PtnCD (IICIID*_L. lactis_*), ManYZ (IICIID*_E. coli_*), and MptCD (IICIID*_L. monocytogenes_*) were compared using the Clustal Omega software at the EMBL-EBI online server (46). Structural elements (HPs, AHs, THs) of PtnCD, ManYZ, and MptCD were taken from the experimentally identified structures available under PDB ID numbers 8HFS, 6K1H, and 7VLX, respectively (3, 22, 23). Structural elements of ManCD were predicted based on the experimentally determined structure of PtnCD. For visualization of PtnCD and ManCD, the Protter server was used (47).

GarQ, GarA, GarB, GarC, and BacSJ sequence homology searches were performed using the BLAST algorithm on the NCBI platform and visualized with MultAlin online software (48). The protein host organisms and accession numbers are listed in Table S1. The bacteriocin amino acid sequences underwent molecular phylogenetic tree inferences using the phylogeny.fr online tool with default settings.

Secondary structures of bacteriocins were predicted using PSIPRED (49) or derived from known tertiary structures deposited in the PDB repository. Tertiary structures were built using I-TASSER web services (50) or derived from PDB and visualized using the PyMOL Molecular Graphics System, Version 2.0 (Schrödinger, LLC). The PDB accession numbers of the templates of the highest significance are: 1OG7A for GarQ, 2YX1A for GarA, 3FD2A for GarC, and 1RY3 for LcnB. The PDB accession numbers for bacteriocins are: Pediocin PA-1—7VLY; Sakacin A—7XNO; Sakacin P— 1OG7; Leucocin A—3LEU; Lactococcin A—8HFS. The amino acid substitutions for the synthesis of GarQ variants, and for site-directed mutagenesis of PtnCD were designed according to the Russelllab web service to ensure their neutral impact on the bacteriocin or mannose transporter structure, respectively (51).

### Site-directed mutagenesis

To express *ptnCD* with the designed mutations, the nisin-controlled gene expression system (NICE) with pNZ9530 and pNZ8037 plasmids was used (52). Recombinant pNZ8037 plasmid carrying wild-type or mutated *ptnCD* genes was isolated using PureYield^TM^ Plasmid Mini-prep System (Promega, USA). Targeted changes in the *ptnCD* genes were introduced following the Site Directed Mutagenesis Cornell iGEM 2012 Protocol (http://2012.igem.org/wiki/images/a/a5/Site_Directed_Mutagenesis.pdf). The pNZ8037:*ptnCD* construct isolated from *E. coli* B529a (7) was used as a template for PCR. To introduce the Ser55Val, Ile59Phe, Val101Phe, Ile105Phe, or Tyr198Val mutation into *ptnC*, mutagenic primer pairs *ptnC*Ser55Valfor/rev, *ptnC*Ile59Phefor/rev, *ptnC*Val101Phefor/rev, *ptnC*Ile105Phefor/rev, or *ptnC*Tyr198Valfor/rev (Table S1), respectively, were used. To introduce the Pro115Val, Thr134Val, or Trp201Val mutation into *ptnD*, mutagenic primer pairs *ptnD*Pro115Valfor/rev, *ptnD*Thr134Valfor/rev, or *ptnD*Trp201Valfor/rev (Table S1), respectively, were used. Multiplied plasmids were transferred into *E. coli* EC1000 and the presence of correct inserts was confirmed by plasmid sequencing with pNZ8037for/rev primer pair (Table S1). Finally, recombinant pNZ8037 plasmids were expressed in *L. lactis* B488.

## Acknowledgements

This work was supported by grant no. 2018/29/N/NZ1/00965 from the National Science Centre (Poland), Foundation for Polish Science (FNP), and Polish National Agency for Academic Exchange (NAWA) as part of the Bekker Scholarship Programme. The authors acknowledge Dr. John Vederas for the opportunity to use the facilities for circular dichroism and peptide mass analysis at the Department of Chemistry, University of Alberta, Canada.

## Data Availability Statement

All data are included in the manuscript or supplementary materials.

