## Supplementary material for "Subclass IId bacteriocins targeting Man-PTS—structural diversity and implications for receptor interaction and antimicrobial activity": Supllemental files

**Table S1. Bacterial strains, plasmids and primers used in this study**

| Strains, plasmids, primers, peptides | Description, NCBI accession number | Source* (reference) |
|---|---|---|
| **Strains** | | |
| *Apilactobacillus kunkeei* AH1 | indicator strain | LMGT NMBU |
| *Apilactobacillus kunkeei* AH38 | indicator strain | LMGT NMBU |
| *Apilactobacillus kunkeei* AH119 | indicator strain | LMGT NMBU |
| *Bacillus cereus* IBB3390 | indicator strain | IBB PAS |
| *Bacillus subtilis* BSB1 | indicator strain | (53) |
| *Campylobacter jejuni* 12 | indicator strain | (54) |
| *Campylobacter jejuni* 480 | indicator strain | (55) |
| *Campylobacter jejuni* 81176 | indicator strain | (56) |
| *Campylobacter coli* 23/1 | indicator strain | (54) |
| *Candida albicans* CAI-4 | indicator strain, | (57) |
| *Carnobacterium maltaromaticum* IBB3447 | indicator strain | IBB PAS |
| *Enterococcus durans* IBB3441 | indicator strain | IBB PAS |
| *Enterococcus faecalis* IBB3439 | indicator strain | IBB PAS |
| *Enterococcus faecalis* IBB3444 | indicator strain | IBB PAS |
| *Enterococcus faecium* LMGT 2783 | indicator strain | LMGT NMBU |
| *Enterococcus faecium* LMGT 2787 | indicator strain | LMGT NMBU |
| *Escherichia coli* EC1000 | indicator strain, host strain | (58) |
| *Escherichia coli* TG1 | indicator strain | (59) |
| *Lacticaseibacillus paracasei* IBB3418 | indicator strain | IBB PAS |
| *Lacticaseibacillus paracasei* IBB3425 | indicator strain | IBB PAS |
| *Lacticaseibacillus paracasei* IBB3426 | indicator strain | IBB PAS |
| *Lacticaseibacillus paracasei* IBB3427 | indicator strain | IBB PAS |
| *Lacticaseibacillus paracasei* IBB3428 | indicator strain | IBB PAS |
| *Lacticaseibacillus paracasei* LOCK 0919 | indicator strain, CP005486.1 | LOCK (60) |
| *Lacticaseibacillus paracasei* subsp. *paracasei* IBB3423 | indicator strain, ASM973948v1 | IBB PAS (61) |
| *Lacticaseibacillus rhamnosus* IBB3429 | indicator strain | IBB PAS |
| *Lacticaseibacillus rhamnosus* LOCK 0900 | indicator strain, CP005484.1 | LOCK (62) |
| *Lacticaseibacillus rhamnosus* LOCK 0908 | indicator strain, CP005485.1 | LOCK (63) |
| *Lacticaseibacillus rhamnosus* GG | indicator strain, AP011548 | Dicoflor (64) |
| *Lactiplantibacillus paraplantarum* IBB3438 | indicator strain | IBB PAS |
| *Lactiplantibacillus plantarum* NC8 | indicator strain, AGRI00000000.1 | LMGT NMBU (65) |
| *Lactiplantibacillus plantarum* WCSF1 | indicator strain, AL935263.2 | LMGT NMBU (66) |
| *Lactiplantibacillus plantarum* IBB3433 | indicator strain | IBB PAS |
| *Lactiplantibacillus plantarum* IBB3436 | indicator strain | IBB PAS |
| *Lactiplantibacillus plantarum* subsp. *plantarum* IBB3434 | indicator strain | IBB PAS |
| *Lactobacillus johnsonii* IBB3155 | indicator strain | IBB PAS |
| *Lactococcus garvieae* IBB3403 | indicator strain | IBB PAS |
| *Lactococcus garvieae* DSM 6783 | indicator strain | DSMZ |
| *Lactococcus garvieae* DSM 20064 | indicator strain | DSMZ |
| *Lactococcus garvieae* DSM 20385 | indicator strain | DSMZ |
| *Lactococcus garvieae* DSM 20684 | indicator strain, ASM244167v1 | DSMZ |

| | | |
|---|---|---|
| *Lactococcus garvieae* DSM 20685 | indicator strain | DSMZ |
| *Lactococcus garvieae* DSM 29394 | indicator strain | DSMZ |
| *Lactococcus garvieae* DSM 100577 | indicator strain | DSMZ |
| *Lactococcus garvieae* IBB66 | indicator strain | IBB PAS |
| *Lactococcus garvieae* IPLA 31405 | indicator strain, ASM26970v1 | IPLA-CSIC (67) |
| *Lactococcus garvieae* PAQ 102015-99 | indicator strain, PAQ99-XT | (68) |
| *Lactococcus lactis* IBB3404 | indicator strain | IBB PAS |
| *Lactococcus lactis* IBB3411 | indicator strain | IBB PAS |
| *Lactococcus lactis* QU5 | indicator strain | LMGT NMBU |
| *Lactococcus. cremoris* IBB3409 | indicator strain | IBB PAS |
| *Lactococcus. lactis* IBB3407 | indicator strain | IBB PAS |
| *Lactococcus lactis* IL1403 | indicator strain, host strain, AE005176 | INRA (69) |
| *Lactococcus raffinolactis* IBB91 | indicator strain | IBB PAS |
| *Leuconostoc lactis* IBB3446 | indicator strain | IBB PAS |
| *Leuconostoc mesenteroides* IBB3442 | indicator strain | IBB PAS |
| *Leuconostoc mesenteroides* IBB3443 | indicator strain | IBB PAS |
| *Ligilactobacillus salivarius* IBB3154 | indicator strain | IBB PAS |
| *Listeria monocytogenes* EGD-e | indicator strain, AL591824.1 | LMGT NMBU (70) |
| *Listeria monocytogenes* ATCC 43526 | indicator strain | ATCC |
| *Listeria monocytogenes* ATCC 15313 | indicator strain | ATCC |
| *Listeria monocytogenes* H7762 | indicator strain | CDC |
| *Pediococcus acidilacti* LMGT 2002 | indicator strain | LMGT NMBU |
| *Pediococcus parvulus* IBB3448 | indicator strain | IBB PAS |
| *Pediococcus pentosaceus* IBB3369 | indicator strain | IBB PAS |
| *Pseudomonas aeruginosa* ATCC 9027 | indicator strain | ATCC |
| *Salmonella typhimurium* TT622 | indicator strain | (71) |
| *Staphylococcus aureus* ATCC 6538 | indicator strain | ATCC |
| *Staphylococcus caprae* DSM-20608 | indicator strain | DSMZ |
| *Staphylococcus delphini* DSM-20771 | indicator strain | DSMZ |
| *Staphylococcus epidermidis* DSM-20044 | indicator strain | DSMZ |
| *Staphylococcus hyicus* DSM-20459 | indicator strain | DSMZ |
| *Staphylococcus intermedius* DSM-20373 | indicator strain | DSMZ |
| *Staphylococcus lugdunesis* DSM-4804 | indicator strain | DSMZ |
| *Staphylococcus pseudintermedius* DSM-21284 | indicator strain | DSMZ |
| *Staphylococcus saprophyticus* DSM-18669 | indicator strain | DSMZ |
| *Staphylococcus schleiferi* DSM-6628 | indicator strain | DSMZ |
| *Streptococcus agalactiae* IBB123 | indicator strain | IBB PAS |
| *Streptococcus agalactiae* IBB130 | indicator strain | IBB PAS |
| *Streptococcus mitis* IBB3449 | indicator strain | IBB PAS |
| *Streptococcus parauberis* IBB272 | indicator strain | IBB PAS |
| *Streptococcus sobrinus* IBB3450 | indicator strain | IBB PAS |
| ***Lactococcus garvieae* IBB3403—spontaneous mutants**** | | |
| LG1-G32, LG34-LG38, LG40 | GarAG3$^{r}$; Man$^{+}$ | This study |
| LGA2, LGA3, LGA6, LGA10 | GarA$^{r}$; Man$^{+}$ | (8) |
| LGB6 | GarB$^{r}$; Man$^{+}$ | (8) |
| LGC1, LGC8, LGC13, LGC15 | GarC$^{r}$; Man$^{+}$ | (8) |
| PW202, PW203, PW204, LGN1, LGN2, LGN9 | GarQ$^{r}$; Man$^{+}$ | (7) |
| ***Lactococcus garvieae* IBB3403—other mutants**** | | |
| B548a | strain with *manABCD* deletion; Man$^{-}$ | (8) |
| ***Lactococcus lactis* IL1403—spontaneous mutants**** | | |
| LLB2, LLB3, LLB17 | RhmB$^{r}$; Man$^{+}$ | This study |
| LLC1, LLC2 | LcnC$^{r}$; Man$^{+}$ | This study |
| LLG1-LLG3, LLG7-LLG9, LLG11, LLG12, LLG14, LLG19, LLG21, LLG25, LLG32, LLG46, LLG50 | EntG1$^{r}$; Man$^{+}$ | This study |
| LLA1, LLA7, LLA8, LLA12, LLA19-LLA22, LLA24, LLA26-LLA30 | MurA$^{r}$; Man$^{+}$ | This study |
| LLN1 | GarQ$^{r}$; Man$^{+}$ | (7) |
| M6, M16, M19, M30 | BacSJ$^{r}$; Man$^{+}$ | (10) |
| ***Lactococcus lactis* IL1403—other mutants**** | | |
| B464 | IL1403 strain with *ptnABCD* deletion; GarABCQ$^{r}$, BacSJ$^{r}$; Man$^{-}$ | LMGT NMBU (6–8) |
| B488 | B464 carrying pNZ9530 | LMGT NMBU (6–8) |
| B520 | B488 carrying pNZ8037; GarABCQ$^{r}$, BacSJ$^{r}$; Man$^{-}$ | LMGT NMBU (6–8) |
| B515 | B488 carrying pNZ8037 with *ptnABCD*; GarAB$^{r}$, GarCQ$^{s}$, BacSJ$^{s}$; Man$^{+}$ | LMGT NMBU (6–8) |
| B538 | B488 carrying pNZ8037 with *ptnC*; GarABCQ$^{r}$, BacSJ$^{r}$; Man$^{-}$ | LMGT NMBU (6–8) |
| B541 | B488 carrying pNZ8037 with *ptnD*; GarABCQ$^{r}$, BacSJ$^{r}$; Man$^{-}$ | LMGT NMBU (6–8) |

| | | |
|---|---|---|
| B529 | B488 carrying pNZ8037 with *ptnCD*; GarAB$^{r}$, GarCQ$^{s}$, BacSJ$^{s}$; Man$^{+}$ | LMGT NMBU (6–8) |
| B557a | B488 carrying pNZ8037 with *manABCD*; GarABCQ$^{s}$, BacSJ$^{r}$; Man$^{+}$ | (8) |
| B558a | B488 carrying pNZ8037 with *manCD*; GarABCQ$^{s}$, BacSJ$^{r}$; Man$^{+}$ | (8) |
| B559a | B488 carrying pNZ8037 with *manC*; GarABCQ$^{r}$, BacSJ$^{r}$; Man$^{-}$ | (8) |
| B560a | B488 carrying pNZ8037 with *manD*; GarABCQ$^{r}$, BacSJ$^{r}$; Man$^{-}$ | (8) |
| B570a | B529 with Ser55Val substitution in *ptnC*; Man$^{+}$ | This study |
| B571a | B529 with Ile59Phe substitution in *ptnC*; Man$^{+}$ | This study |
| B572a | B529 with Val101Phe substitution in *ptnC*; Man$^{+}$ | This study |
| B573a | B529 with Ile105Phe substitution in *ptnC*; Man$^{+}$ | This study |
| B574a | B529 with Tyr198Val substitution in *ptnC*; Man$^{+}$ | This study |
| B575a | B529 with Pro115Val substitution in *ptnD*; Man$^{+}$ | This study |
| B576a | B529 with Thr134Val substitution in *ptnD*; Man$^{+}$ | This study |
| B577a | B529 with Trp201Val substitution in *ptnD*; Man$^{+}$ | This study |
| ***Escherichia coli* EC1000** | | |
| B529a | strain with pNZ8037 harboring *ptnCD* | (7) |
| B570a | B529a with Ser55Val substitution in *ptnC*; Man$^{+}$ | This study |
| B571a | B529a with Ile59Phe substitution in *ptnC*; Man$^{+}$ | This study |
| B572a | B529a with Val101Phe substitution in *ptnC*; Man$^{+}$ | This study |
| B573a | B529a with Ile105Phe substitution in *ptnC*; Man$^{+}$ | This study |
| B574a | B529a with Tyr198Val substitution in *ptnC*; Man$^{+}$ | This study |
| B575a | B529a with Pro115Val substitution in *ptnD*; Man$^{+}$ | This study |
| B576a | B529a with Thr134Val substitution in *ptnD*; Man$^{+}$ | This study |
| B577a | B529a with Trp201Val substitution in *ptnD*; Man$^{+}$ | This study |
| **Plasmids** | | |
| pNZ9530 | Em$^{r}$, carrying nisin-regulatory *nisRK* genes | (72) |
| pNZ8037 | Cam$^{r}$, nisin regulated expression system with nisin-responsive promoter | (52) |
| **Primers** | Nucleotide sequence (5'->3') | |
| *ptnC*for/rev | TCTGACCTCTTTGGTTTG/GCATACGTTCGTAGT | |
| *ptnD*for/rev | AACCTCCAAGCTTCTG/AGCCACAGATTCTCTCC | |
| *manC*for/rev | CGTGATCTCGGCGTTA/TAACGCTCAAGCGTGTG | |
| *manD*for/rev | CGCTCTTATCTACCTC/GCCAATTTAGTGCTCCTAAC | |
| pNZ8037for/rev | CGATAACGCGAGCATA/GCTCAAGGGCTTTTACG | |
| *ptnC*Ser55Valfor/rev | GCAGGGATTATCCTCGGTGGTGTTCTTCAATTGATCGCTCTTGGT/<br>ACCAAGAGCGATCAATTGAAGAACACCACCGAGGATAATCCCTGC | |
| *ptnC*Ile59Phefor/rev | CGGTGGTTCACTTCAATTGTTTGCTCTTGGTTGGGCTAACG/<br>CGTTAGCCCAACCAAGAGCAAACAATTGAAGTGAACCACCG | |
| *ptnC*Val101Phefor/rev | CACATCATGGGTACTATCTTTCCTGCTGCTATCTTGC/<br>GCAAGATAGCAGCAGGAAAGATAGTACCCATGATGTG | |
| *ptnC*Ile105Phefor/rev | TACTATCGTTCCTGCTGCTTTTTTGCTTGCAACTGCTGGTC/<br>GACCAGCAGTTGCAAGCAAAAAAGCAGCAGGAACGATAGTA | |
| *ptnC*Tyr198Valfor/rev | TGGGATGGTTGTTGCCGTTGGTGTTGCAATGGTTATCAACC/<br>GGTTGATAACCATTGCAACACCAACGGCAACAACCATCCCA | |
| *ptnD*Pro115Valfor/rev | CTTGCCGGTATCGGTGACGTTGTCTTCTGGTTTACAGT/<br>ACTGTAAACCAGAAGACAACGTCACCGATACCGGCAAG | |
| *ptnD*Thr134Valfor/rev | TGGTGCGATTGCAGCTTCATTGGCTGTTGGTGGATCAATT/<br>AATTGATCCACCAACAGCCAATGAAGCTGCAATCGCACCA | |
| *ptnD*Trp201Valfor/rev | TTCGTCCTTGGTGTATTGATTCAACGTGTTGTAACAATTAACTTTAATGGTCCTAAC/<br>GTTAGGACCATTAAAGTTAATTGTTACAACACGTTGAATCAATACACCAAGGACGAA | |
| **Peptides** | Host organism | NCBI accession number(s) |
| GarQ | *Lactococcus garvieae* BCC 43578, plasmid | AEN79392.1 |
| GarAG2 | *Lactococcus garvieae* Lg-Granada, plasmid | WP_165719065.1;<br>NZ_CP084378.1 |
| LcnC/HB1 | *Lactococcus cremoris* UC073 | UXV60242.1 |
| HB2 | *Lactococcus garvieae* FDAARGOS_1002, plasmid | QQB43181.1 |
| EntG1/HB3 | *Enterococcus* sp. HSIEG1 | EQC79567.1 |
| HB4 | *Ligilactobacillus agilis* 268A | PLA76810.1 |
| AglA/HB5 | *Ligilactobacillus agilis* UMB7763 | MDK6810183.1 |
| HB6 | *Ligilactobacillus agilis* C7 | MCL8204652.1 |
| MurA/HB7 | *Ligilactobacillus murinus* DSM 100193 | MCR1897247.1 |
| HB8 | *Lactobacillus equicursoris* DSM 19284 | CCK84999.1 |
| HB9 | *Furfurilactobacillus rossiae* DSM 15814 | KRL52547.1 |
| Bov255 | *Streptococcus* sp. LRC 0255 | AAG29818.1 |
| Ubericin K | *Streptococcus uberis* LMGT 4214 | QWX09803.1 |
| Angicin | *Streptococcus anginosus* BSU 1211 | UJH61018.1 |
| HB10 | *Leuconostoc carnosum* WC0328 | KAA8327172.1 |
| HB11 | *Lacticaseibacillus paracasei* N53 | MBU5324208.1 |

| | | |
|---|---|---|
| RhmB/HB12 | *Lacticaseibacillus rhamnosus* VHProbi M14 | UUT37922.1 |
| HB13 | *Lacticaseibacillus rhamnosus* TOM.283 | MDM7524757.1 |
| BacSJ | *Lacticaseibacillus paracasei* subsp. *paracasei* BGSJ2-8, plasmid | CAR92206.2 |
| LcbC/HB14 | *Companilactobacillus kimchii* DSM 13961 | KAE9557329.1 |
| HB15 | *Lactococcus lactis* Bpl1 | KGF75926.1 |
| HB16 | *Lactococcus lactis* 1031 | MCT1192459.1 |
| LcnA | *Lactococcus cremoris* 9B4, plasmid | P0A313.1 |
| HB17 | *Lactococcus lactis* FDAARGOS_866 | WP_228777672.1, NZ_CP065734.1 |
| LcnZ | *Lactococcus lactis* QU7 | BAU29928.1 |
| LcnB | *Lactococcus cremoris* 9B4, plasmid | P35518.1 |
| HB18 | *Lactococcus cremoris* TIFN3 | EQC95783.1 |
| HB19 | *Lactobacillus* sp. CBA3605 | AVK61836.1 |
| HB20 | *Lactococcus lactis* 537 | MDG4972525.1 |
| GarC | *Lactococcus garvieae* 21881, plasmid | CCF55362.1 |
| LcnD/HB21 | *Lactococcus* sp. | MBR6895708.1 |
| HB22 | *Enterococcus gallinarum* ENT.1 | MBF0724730.1 |
| MsnC/HB23 | *Leuconostoc mesenteroides* 213M0 | WP_061399677.1, NZ_BCMO01000030.1 |
| HB24 | *Leuconostoc mesenteroides* INF3b | MBZ1529843.1 |
| HB25 | *Leuconostoc mesenteroides* subsp. m*esenteroides* FM06, plasmid | ARR90014.1 |
| HB26 | *Leuconostoc citreum* LBAE C11 | CCF27630.1 |
| HB27 | *Lactococcus lactis* 537 | PFG87625.1 |
| HB28 | *Lactococcus* sp. EKM201L | KAF6605648.1 |
| HB29 | *Lactococcus cremoris* B40 | KZK48428.1 |
| HB30 | *Lactococcus lactis* | MEE0824874.1 |
| HB31 | *Enterococcus faecalis* TX4248 | EFM82101.1 |
| GarA | *Lactococcus garvieae* 21881, plasmid | CCF71073.1 |
| GarB | *Lactococcus garvieae* 21881, plasmid | CCF55365.1 |
| GarAG3/HB32 | *Lactococcus garvieae* M14, plasmid | CEF52411.1 |
| GarAG1 | *Lactococcus garvieae* Lg-Granada, plasmid | WP_225667055.1, NZ_CP084378.1 |

*Strains derive from the collections of: the Laboratory of Microbial Gene Technology, Department of Chemistry, Biotechnology and Food Science, Norwegian University of Life Sciences, Ås, Norway (LMGT NMBU); the Regional Strains and Plasmids Collection of the Institute of Biochemistry and Biophysics, Warsaw, Poland (IBB PAS); the Pure Cultures Collection of the Institute of Fermentation Technology and Microbiology, Technical University of Lodz, Poland (LOCK); the dietary supplement Dicoflor, Vitis Pharma, Poland (Dicoflor); the Collection of Microorganisms and Cell Cultures, Germany (DSMZ); the collection of the Dairy Institute of Asturias, Spanish National Research Council, Asturias, Spain (IPLA-CSIC); the collection of the National Institute for Agricultural Research, France (INRA), the American Type Culture Collection (ATCC), Centers for Disease Control and Prevention (CDC), otherwise they were obtained in earlier studies or in this study.

** Man – mannose phenotype (ability to grow on mannose as a sole carbon source)

**Table S2. Sensitivity of *L. garvieae* IBB3403 and *L. lactis* IL1403 mutants to peptides with similarity to Man-PTS bacteriocins**

| Indicator strain | Genotype | MurA | MsnC | RhmB | AglA | LcbC | LcnC | LcnD | GarAG3 | EntG1 | HB10 HB13 HB15 HB19 HB20 HB27 |
|---|---|---|---|---|---|---|---|---|---|---|---|
| *L. garvieae* | | | | | | | | | | | |
| IBB3403 | wild-type | + | +/- | - | - | - | ++ | + | + | - | - |
| B548a | Δ*manABCD* | - | - | - | - | - | - | - | - | - | - |
| *L. lactis* | | | | | | | | | | | |
| IL1403 | wild-type | + | +/- | + | + | + | ++ | + | - | + | - |
| B464 | Δ*ptnABCD* | - | - | - | - | - | - | - | - | - | - |
| *L. lactis* Δ*ptnABCD* | | | | | | | | | | | |
| B520 | pNZ8037 | - | - | - | - | - | - | - | - | - | - |
| B557a | p*manABCD* | + | +/- | - | - | - | ++ | + | + | - | - |
| B558a | p*manCD* | + | +/- | - | - | - | ++ | + | + | - | - |
| B559a | p*manC* | - | - | - | - | - | - | - | - | - | - |
| B560a | p*manD* | - | - | - | - | - | - | - | - | - | - |
| B515 | p*ptnABCD* | + | +/- | + | + | + | ++ | + | - | + | - |
| B529 | p*ptnCD* | + | +/- | + | + | + | ++ | + | - | + | - |
| B538 | p*ptnC* | - | - | - | - | - | - | - | - | - | - |
| B541 | p*ptnD* | - | - | - | - | - | - | - | - | - | - |

5 µl of each bacteriocin at a concentration of 1 mg/ml was used; "–", no inhibition zone (strain resistance); "+/–", minimal, vague inhibition zone (minimal strain sensitivity); "+" and "++", wide, clear inhibition zone with, respectively, diameter ≤ 10 mm and >10 mm (strain sensitivity)

**Table S3. Spontaneous mutants of *L. lactis* IL1403 resistant to RhmB, EntG1, LcnC, MurA, GarQ, BacSJ, ubericin K, and GarC**

| Mutant strain | Mutation | Amino acid change | Sensitivity to [fold-decreased relative to WT] | | | | | | | | Position in the cell membrane of affected Man-PTS amino acid(s) |
|---|---|---|---|---|---|---|---|---|---|---|---|
| | | | RhmB | EntG1 | LcnC | MurA | GarQ | BacSJ | Ubericin K | GarC | |
| RhmB-resistant mutants | | | | | | | | | | | |
| LLB2,LLB3 | *ptnD*: 376G>A | PtnD: Gly126Ser | 8x | 4x | 0x | 32x | 0x | >4x | 4x | 0x | transmembrane (TH1) |
| LLB17 | *ptnD*: 641C>T | PtnD: Ser214Leu | >16x | >8x | >128x | 256x | 256x | >4x | 32x | 0x | outside |
| EntG1-resistant mutants | | | | | | | | | | | |
| LLG1, LLG2 | *ptnD*: 531-681multiple mutations | PtnD: 221-228del GlyAlaTyrLeuGluPheProLys | >16x | >8x | 16x | >1024x | 16x | >4x | >128x | >32x | outside (region γ) |
| LLG3, LLG9, LLG11, LLG32, LLG46 | *ptnD*: 686G>T | PtnD: Gly229Val | >16x | >8x | 16x | 8x | 16x | >4x | 0x | >32x | outside (region γ) |
| LLG7, LLG19, LLG21 | *ptnD*: 379G>A | PtnD: Ala127Thr | >16x | >8x | 8x | 16x | 8x | >4x | 32x | >32x | transmembrane (TH1) |
| LLG8 | *ptnD*: 661G>T | PtnD: Gly221Cys | 8x | >8x | 16x | 4x | 16x | >4x | 0x | >32x | outside |
| LLG12, LLG14 | *ptnC*: 560C>T | PtnC: Ala187Val | 0x | >8x | 8x | 32x | 8x | >4x | 2x | >32x | transmembrane (TH3) |
| LLG25 | *ptnD*: 668A>G | PtnD: Tyr223Cys | >16x | >8x | 32x | 64x | 16x | >4x | 2x | >32x | outside (region γ) |
| LLG50 | *ptnC*: 360G>T | PtnC: Met120Ile | 8x | >8x | 4x | 64x | 4x | >4x | 8x | 8x | transmembrane (TH1) |
| LcnC-resistant mutants | | | | | | | | | | | |
| LLC1, LLC2 | *ptnC*: 185G> T | PtnC: Gly62Val | >16x | >8x | 4x | 128x | 64x | >8x | 64x | >32x | transmembrane |
| MurA-resistant mutants | | | | | | | | | | | |
| LLA1, LLA7, LLA8, LLA12, LLA19-LLA22, LLA24, LLA26-LLA30 | *ptnC*: 328G>A | PtnC: Ala110Thr | 4x | >8x | 2x | >1024x | 4x | >4x | 64x | 0x | transmembrane (TH1) |
| GarQ-resistant mutants | | | | | | | | | | | |
| LLN1 | *ptnD*: 368C>A | PtnD: Pro123His | >16x | >8x | >128x | 64x | >1024x | >8x | >128x | 8x | transmembrane (TH1) |
| BacSJ-resistant mutants | | | | | | | | | | | |
| M6 | *ptnD*: 599G>A | PtnD: Arg200His | 8x | 4x | 2x | 4x | 8x | >8x | 2x | >32x | transmembrane (TH3) |
| M16 | *ptnD*: 247C>T | PtnD: Leu83Phe | 8x | 4x | 2x | 8x | 0x | >8x | 4x | 4x | transmembrane (HP2b) |
| M19 | *ptnD*: 677T>C | PtnD: Phe226Ser | >16x | >8x | 8x | 16x | 8x | >8x | 0x | >32x | outside (region γ) |
| M30 | *ptnD*: 591G>T | PtnD: Leu197Phe | >16x | >8x | 16x | 32x | 16x | >8x | 4x | >32x | transmembrane (TH3) |

“>” substitution; “del”deletion

**Table S4. Spontaneous mutants of *L. garvieae* IBB3403 resistant to GarAG3, GarA, GarB, GarC, GarQ, MurA, and ubericin K**

| Mutant strain | Mutation | Amino acid change | Sensitivity to [fold-decreased relative to WT] | | | | | | | Position in the cell membrane of affected Man-PTS amino acid(s) |
|---|---|---|---|---|---|---|---|---|---|---|
| | | | GarAG3 | GarA | GarB | GarC | GarQ | MurA | Ubericin K | |
| GarAG3-resistant mutants | | | | | | | | | | |
| LG1, 3-5, 7, 8, 10, 14-17, 19-21, 24-27, 30, 31, 34, 35, 37, 38, 40 | *manD*: 608G >T | ManD: Arg203Leu | >1024x | 0x | >64x | >128x | 0x | 0x | 4x | transmembrane (TH3) |
| LG2 | *manC*: 178G>T | ManC: Gly60Cys | 1024x | 4x | >64x | >128x | 32x | >16x | >32x | transmembrane |
| LG6 | *manD*: 767G>T | ManD: Trp256Leu | 128x | 0x | 4x | 8x | 16x | 0.5x | 4x | extracellular (region γ+) |
| LG9 | *manD*: 800G>T | ManD: Gly267Val | 16x | 0x | 8x | 2x | 4x | 0.5x | 4x | extracellular (region γ+) |
| LG11, 13, 22, 28 | *manD*: 607C>T | ManD: Arg203Cys | >1024x | 0x | >64x | >128x | 2x | 0x | 2x | transmembrane (TH3) |
| LG12 | *manD*: 788_799del TTGTTGGTAACG | ManD: 262_265delAsnValValGly | 64x | 2x | >64x | 8x | 4x | 0.25x | 0x | extracellular (region γ+) |
| LG18 | *manD*: 794_805del GTAACGGTGTTG | ManD: 265_268delGlyAsnGlyVal | 32x | 2x | 8x | 8x | 4x | 0.0625x | 2x | extracellular (region γ+) |
| LG23, 29 | *manC*: 563G>T | ManC: Gly188Val | >1024x | 2x | >64x | >128x | 4x | 0x | 8x | transmembrane (TH3) |
| LG32 | *manD*: 953C>T | ManD: Ser318Leu | 4x | 2x | 32x | >128x | 2x | 0x | >32x | extracellular |
| LG36 | *manC*: 179G>T | ManC: Gly60Val | >1024x | 8x | >64x | >128x | 128x | >16x | >32x | transmembrane |
| GarA-resistant mutants | | | | | | | | | | |
| LGA2 | *manC*: 176T>C | ManC: Leu59Pro | 32x | 32x | 32x | 32x | 128x | 4x | >32x | transmembrane (HP2a) |
| LGA3 | *manC*: 155G>T | ManC: Gly52Val | 32x | 4x | 64x | 16x | 4x | 0.5x | 8x | transmembrane (HP2a) |
| LGA6 | *manC*: 570_571insGTT | ManC:190_191insVal | 128x | 8x | 32x | 64x | 4x | 0.5x | 0x | transmembrane (TH3) |
| LGA10 | *manD*: 173C>A | ManD: Ala58Asp | 8x | 2x | 4x | 4x | 2x | 2x | 8x | transmembrane (HP2a) |
| GarB-resistant mutant | | | | | | | | | | |
| LGB6 | *manD*: 937C>A | ManD: Gln313Lys | 2x | 0x | >64x | 2x | 2x | 0.25x | 4x | extracellular |
| GarC-resistant mutants | | | | | | | | | | |
| LGC1 | *manD*: 102G>A | ManD: Met34Ile | 8x | 0x | 0x | >128x | 0x | 0x | 16x | transmembrane (HP1b) |
| LGC8 | *manD*: 676T>G | ManD: Tyr226Asp | >1024x | 8x | >64x | >128x | 16x | 0x | 0x | extracellular (region γ) |
| LGC13 | *manC*: 317C>T | ManC: Ala106Val | >1024x | 2x | >64x | >128x | 4x | 4x | >32x | transmembrane (TH1) |
| LGC15 | *manD*: 398C>T, 943G>A | ManD: Ala133Val, Val315Met | 32x | 2x | >64x | >128x | 4x | 0x | 8x | transmembrane (TH1), extracellular |
| GarQ-resistant mutants | | | | | | | | | | |
| PW202 | *manC*: 299C>A | ManC: Pro100His | 2x | 4x | 0x | 8x | >1024x | >16x | >32x | transmembrane (TH1) |
| PW203 | *manD*: 368C>T | ManD: Thr123Ile | 64x | 0x | >64x | 0x | >1024x | 0x | >32x | transmembrane (TH1) |
| PW204 | *manD*: 331C>T | ManD: Pro111Ser | 64x | 0x | >64x | 0x | >1024x | >16x | >32x | transmembrane (TH1) |
| LGN1 | *manD*: 902G>T | ManD: Gly301Val | 256x | 8x | >64x | 32x | >1024x | 2x | 0x | extracellular (region γ) |
| LGN2 | *manD*: 398C>T | ManD: Ala133Val | 2x | 0x | 4x | 16x | 16x | 2x | >32x | transmembrane (TH1) |
| LGN9 | *manC*: 305C>T | ManC: Ala102Val | 64x | 0x | >64x | 8x | 16x | 0.125x | 0x | transmembrane (TH1) |

5 µl of each bacteriocin at a concentration of 1 mg/ml was used; “>” substitution; “del”deletion; “ins” insertion

**Table S5.** ***L. lactis*** **B529 strains with site-specific mutations resistant to selected bacteriocins**

| Mutant strain (Amino acid change) | Diameter of the inhibition zone [mm] | | | | | | | | | | | | |
|---|---|---|---|---|---|---|---|---|---|---|---|---|---|
| | GarQ | BacSJ | GarC | Ubericin K | MurA | MsnC | Angicin | RhmB | AglA | LcbC | LcnC | LcnD | EntG1 |
| B520 | - | - | - | - | - | - | - | - | - | - | - | - | - |
| B529 | 24 | 12 | +/- | 16 | 9 | +/- | 7 | +/- | 9 | 7 | 19 | 6 | +/- |
| B570a (Ser55Val in PtnC) | 9 | +/- | - | 11 | - | - | - | - | - | - | 12 | - | - |
| B571a (Ile59Phe in PtnC) | 8 | - | - | - | - | - | - | - | - | - | +/- | - | - |
| B572a (Val101Phe in PtnC) | 10 | - | - | +/- | +/- | - | - | - | - | - | 19 | - | - |
| B573a (Ile105Phe in PtnC) | 10 | - | - | +/- | +/- | - | - | - | 7 | - | 13 | 6 | - |
| B574a (Tyr198Val in PtnC) | 9 | - | - | 10 | +/- | - | - | - | - | - | 12 | - | - |
| B575a (Pro115Val in PtnD) | 10 | - | - | +/- | +/- | - | - | - | - | - | 13 | - | - |
| B576a (Thr134Val in PtnD) | 10 | - | - | 12 | +/- | - | - | - | +/- | - | 14 | - | - |
| B577a (Trp201Val in PtnD) | 10 | - | - | 15 | 9 | - | - | - | +/- | - | 15 | 6 | - |

"–", no inhibition zone (strain resistance); "+/–", minimal, vague inhibition zone (minimal strain sensitivity)

**Figure S1. Phylogenetic tree of peptides with similarity to Man-PTS-binding bacteriocins.** The scale bar shows the evolutionary distance. Bootstrap values (%) are given in nodes.

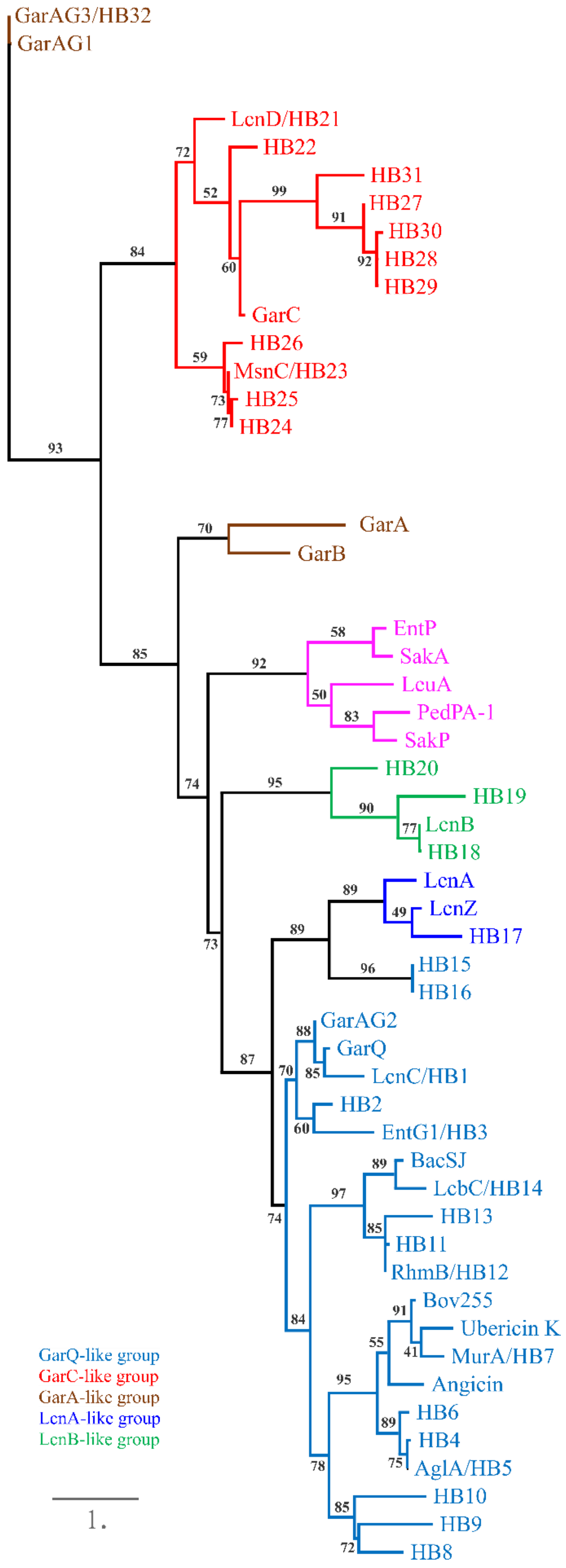

**Figure S2. Tertiary structures of Man-PTS-binding bacteriocins.** GarQ, GarA, GarC, and LcnB structures were predicted *in silico*, and PedPA-1 and LcnA structures were derived from PDB repository. The confidence in predicting tertiary structure is represented by a confidence score (C-score). The C-score typically ranges from -5 to 2, with a higher C-score indicating a more reliable model (50) . Localization of amino acid motifs conserved within the GarQ- and PedPA-1-like group is indicated.

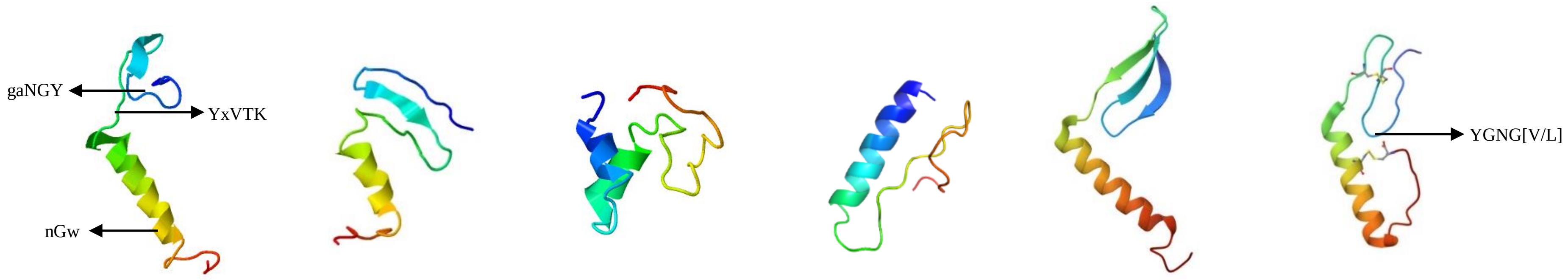


GarQ C-score=-4.59 | LcnB C-score=-5 | GarC C-score=-5 | GarA C-score=-2.16 | LcnA PDB acc. no.: 8HFS | PedPA-1 PDB acc. no.: 7VLY

**Figure S3. Inhibitory spectrum of peptides with similarity to Man-PTS-binding bacteriocins and of mutated variants of GarQ.** Activity of GarQ, GarA, GarB, GarC, and BacSJ was partially taken from our earlier studies (7, 8, 10). Activity of LcnA and LcnB was predicted based on literature data (6). “no activity”, no inhibition zone (strain resistance); “low activity”, minimal, vague inhibition zone (minimal strain sensitivity); “medium activity” and “high activity”, wide, clear inhibition zone with, respectively, diameter ≤ 10 mm and >10 mm (strain sensitivity). GarQ-, GarC-, GarA-, LcnA-, and LcnB-like bacteriocins are in blue, red, brown, navy, and green, respectively. The presence of Man-PTS region(s) α, γ, and γ+ is indicated in the third column. G− and G+ indicate Gram-negative and Gram-positive bacteria, respectively. *, does not contain the GG site of protease processing, thus no signal peptide was removed.

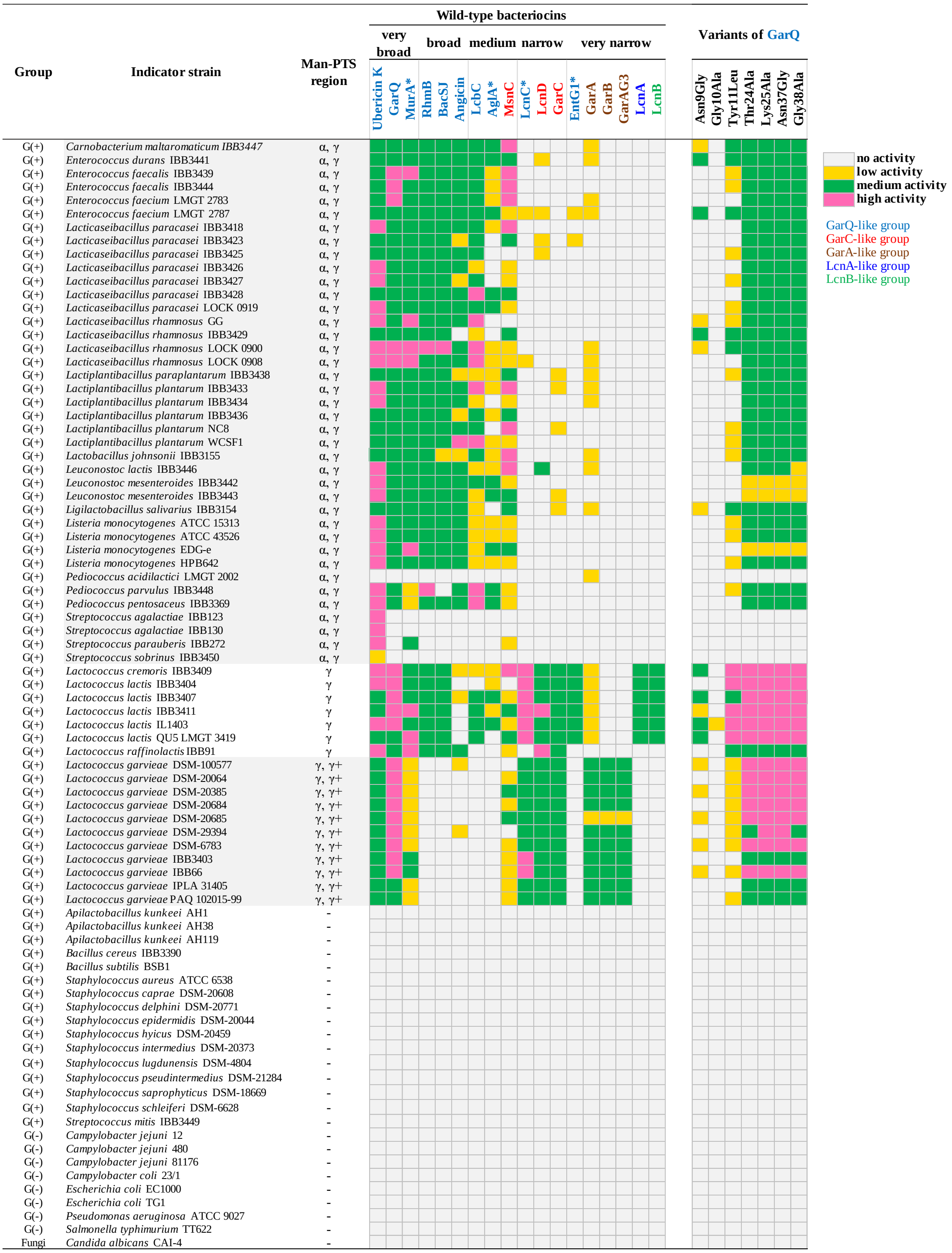

**Figure S4. CD spectra of the HB10, HB13, HB15, HB19, HB20 and HB27 peptides with no antimicrobial activity.** *, does not contain the GG site of protease processing, thus no signal peptide was removed.

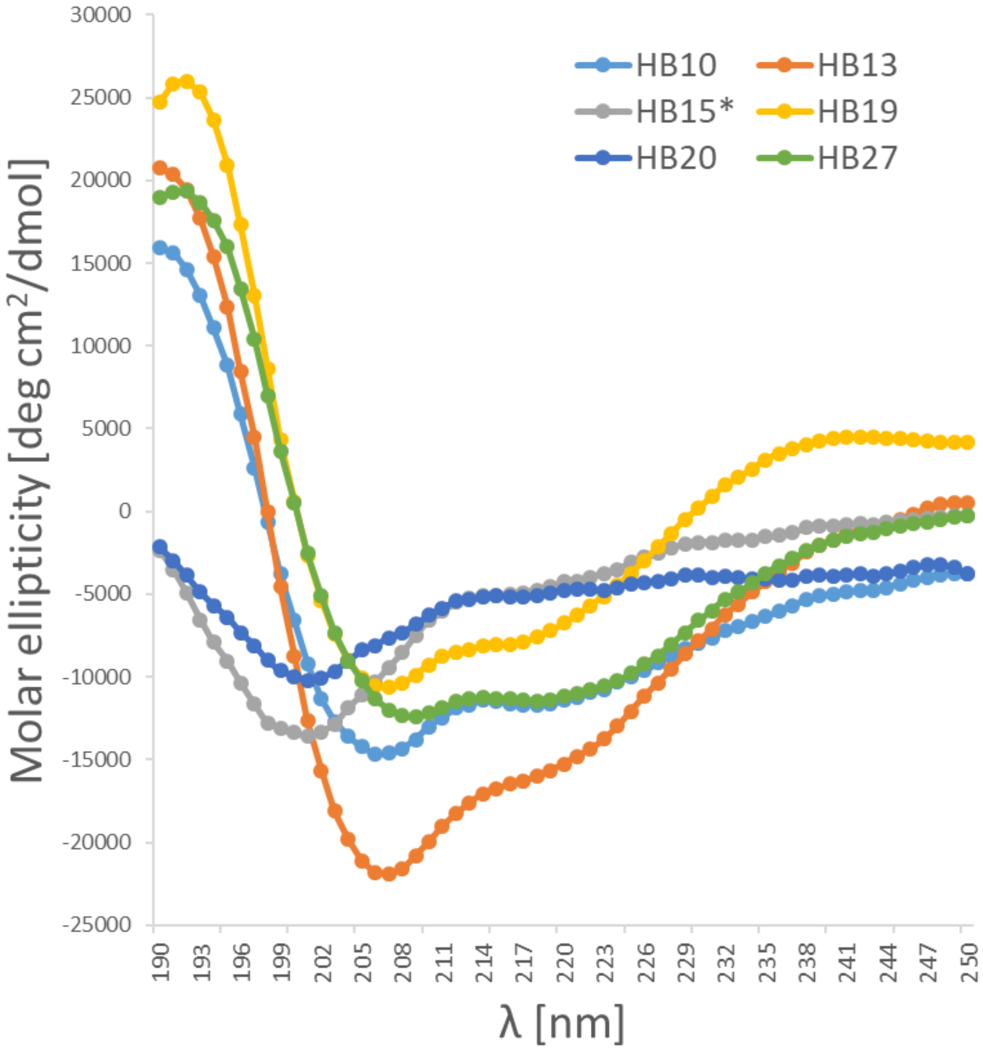

**Figure S5. Sequence alignment of Man-PTS subunits IICIID from *E. coli*, *L. monocytogenes*, *L. lactis*, and *L. garvieae*.** Asterisks, colons, and dots indicate fully, strongly and weakly conserved residues, respectively. Structural element symbols are: HP—hairpin, AH—amphipathic helix, TH—transmembrane helix. Within each subunit, coloured background was used to indicate: Core domain formed by HPs, TH1 and TH2—orange, Vmotif domain formed by TH3-TH5—blue, and AH—pink. The α, β, γ, and γ+ regions are underlined. The residues of Man-PTS$_{E.\ coli}$ involved in mannose or MccE492 binding are indicated by red or black background, respectively (3, 27). The residues of Man-PTS$_{L.\ monocytogenes}$ involved in interaction with PedPA-1 are indicated by grey background (22). The residues of Man-PTS$_{L.\ lactis/L.\ garvieae}$ affected by spontaneous or targeted mutations in bacteriocin-resistant mutants are indicated by green and navy blue background, respectively. Conserved residues changed by the mutations are in green boxes.

### IIC

```
                               HP1a                     HP1b             HP2a                         HP2b      region α             TH1
E. coli           MEITTLQIVLVFIVACIAGMGSILDEFQFHRPLIACTLVGIVLGDMKTGIIIGGTLEMIALGWMNIGAAVAPDAALASIISTILVIAGHQS-----IGAGIALAIPLAAAGQVLTIIVRTITV  118
L. monocytogenes  --MSVISIILVVLIAFLAGIEGILDEFQFHQPLIACTLIGLVTGNLTACIILGGTLQMIALGWANIGAAVAPDAALASVASAIILVLGGQGVAG-I-PSAIAIAIPLAVAGLFLTMIVRTLAV  119
L. lactis         MEYGVLSVILVIVVAFLAGLEGILDQWQFHQPIIACSLIGIVTGHASAGIILGGSLQLIALGWANVGAAVAPDAALASIASSILMVQSNNFDLTHIMGTIVPAAILLATAGLVLTTLVRMLSV  123
L. garvieae       --MSIISIILVLIFAFLAGLEGILDQWQFHQPIIACSLIGIATGHMAAGIILGGSLQMIALGWANVGAAVAPDAALASVASAILMVQGGNFDLTHITGVIVPAAILLATAGLVLTTLVRFLSV  121
                        :.::**.:.*  :**:  .***::***:*:***:*:*:.  *.   :  **:**:*::*****  *:************:  *:*:::  .  :              :   **  **.**  .**  :**  ::*

                                             TH2                 AH              TH3                       TH4                  TH5
E. coli           AFQHAADKAADNGNLTAISWIHVSSLFLQAMRVAIPAVIVALSVGTSEVQNMLNAIPEVVTNGLNIAGGMIVVVGYAMVINMMRAGYLMPFFYLGFVTAAFTNFNLVALGVIGTVMAVLYIQL  241
L. monocytogenes  PIVHLMDRAAEKGNIRSVEWLHISAICMQGIRIAIPAAAL-LFIPADSVQSFLEAMPAWLTDGMAIGGGMVVAVGYALVINMMATKEVWPFFVIGFVVAAISQLTLIAIGALGVALALIYLNL  241
L. lactis         VLVHQADRAAENGSYSGVEMWHFIALICQGLRIAIPAGLL-LVISPDAIQKALAAIPPVISGGLAVGGGMVVAVGYAMVINLMATREVWPFFFLGFALAPISELTLIATGVLGVVIAIVYLNL  245
L. garvieae       GIVHLADAAAEKGSYSGVAGWHMFALLLQGLRIAIPAAII-LAIPAETVTAALNAIPDWVSKGLAVGGGMVVVVGYAMVINLMATKELWPFFFLGFVLAPLSSITLIGMGILGVVIALIYLNL  243
                    :  *   * **::*.   .:    *. ::    *.:*:****   :  * :   .  :     *  *:*    :: *:  :.***:*.****:***:*  :   :  ***  :**.  *  ::.:.*:.  *  :*..:*::*::*

                                region β
E. coli           SPKYNRVAGAPAQAAGNNDLDNELD--   266
L. monocytogenes  SKMGGGNSNGGGGGNSRDPLGDILNDY   268
L. lactis         QASGGSG--NGTASSSGDPIGDILNDY   270
L. garvieae       SNTAGNG---GTASSSGDPIGDILNDY   267
                  .   .         .  . :  :.:  *:
```

### IID

```
                                   HP1a                        HP1b              HP2a                    HP2b                                TH1
E. coli           MVDTTQTTTEKKLTQSDIRGVFLRSNLFQGSWNFERMQALGFCFSMVPAIRRLYPENNEARKQAIRRHLEFFNTQPFVAAPILGVTLALEEQRANGAEIDDGAINGIKVGLMGPLAGVGDPIF  123
L. monocytogenes  ------MAEKIELSKRDRLRVAWRSTFIQGSWNYERMQNGGWAFSMIPAIKKLYKTK-EDRSSALKRHLEFFNTHPYIASPILGVTLALEEERANGAEVDDVAIQGVKVGMMGPLAGVGDPVF  116
L. lactis         -----MSENKVTLDKKIRRSVMWRSMFLQGSWNYERMQNGGWAYSLIPALKKLYPSG-EEAKEALKRHLEFFNTHPYVAAPIIGVTLALEEERANGADIDDAAIQGVKVGMMGPLAGIGDPVF  117
L. garvieae       ---MENTNKKFSLSKNTRLSVMFRSMFLQGSWNYERMQNLGFLYSIIPALKQFYKPGSEEAKEALKRHMEFFNTHPYVAAPIVGVTLALEEEIANGVEIDEAAIQGVKVGMMGPLAGIGDPVF  120
                            :  * :      *   **  ::*****:****   *:  :*::**::::*     *   ..*::**:*****:*::*:**:********:  ***.::*:  **:*:***:******:***:*

                                                       TH2             AH          TH3
E. coli           WGTVRPVFAALGAGIAMSGSLLGPLLFFILFNLVRLATRYYGVAYGYSKGIDIVKDMGGGFLQKLTEGASILGLFVMGALVNKWTHVNIPLVV---SRITD----------------------  221
L. monocytogenes  WFTIRPMLGALGASLALSGNILGPILFFVAWNVIRWGFMWYTQEFGYKAGSKITDDLSGGLLQDITKGASILGMFVLAALVQRWVNIQFAP---IISKVKLDEGAYIDWSHLPQG--------  228
L. lactis         WFTVRPIVGAIAASLATGGSIIAPLFFFIVWNAIRIAFLWYTQEFGYKSGSAITKDLGGGLLQTVTKGASILGMFVLGVLIQRWVTINFNGPNAVVSKIPLQKGAYLEFP-------------  227
L. garvieae       WFTVRPIVGAIAASLATGGSIIAPIFFFVVWNAIRIAFLWYTQEFGYKQGTAITSDLGGGMLQQITKGASILGMFILGVLIQRWVNISFTGPNAMLPSKPLADGAYVGEWIDKAGKVVVQGAQ  243
                  * *:**:..*:.*.:*  .*.::.*::**:  :*  :*  .   :*    :**.  *   *..*:.**:**  :*:******:*::..*:::*.  :.:

                                   region γ and region γ+                                                  TH4                    TH5
E. coli           -----------------------------------------------------------QTGKEHVTTVQTILDQLMPGLVPLLLTFACMWLLRK---KVNPLWIIVGFFVIGIAGYACGLLGL 283
L. monocytogenes  --------------------------------------------AQGIKTALEQQQAGLALSEIKVTTLQNNLDNLIPGLAAVALTFLCMWLLKK---KISPIIIILGLFVVGIVGHLIGLL-- 303
L. lactis         ---------------------------------------KGSVSGTQLHDILGQVGNKLSLDPTKVTYLQDNLNQLIPGLAGLLITFLCMWLLKK---KVSPIVIIFGLFVVGILGRWAQIM-- 307
L. garvieae       TGTTGDGVAKFDWLDQAGNVVGNGVAGQGGFAHYVTVDQLNTVDGSTLHNILGQVSSGLGLSPEQTQSLQDVFNSLIPGFIALLLTFLVLWILRKWKNKNAPLFIIIGMFVLGIVLHVAGLA-- 365
                                                                                            .   :.   :*   ::.*:**:   :  :**   :*:*:*     *   *:  **.*:**:**       :
```

**Figure S6. Predicted structural elements of the membrane Man-PTS subunits IIC and IID of *L. lactis* IL1403 (A) and *L. garvieae* IBB3403 (B).** Structural element symbols are: HP—hairpin, TH—transmembrane helix. HPs, TH1, and TH2 of IICIID subunits form the Core domain, and TH3-TH5 of IICIID subunits form the Vmotif domain. Spontaneous mutants obtained in the presence of GarQ, GarA, GarB, GarC, or BacSJ and their levels of resistance to these bacteriocins are from refs. (7, 8, 10).

**(A)**

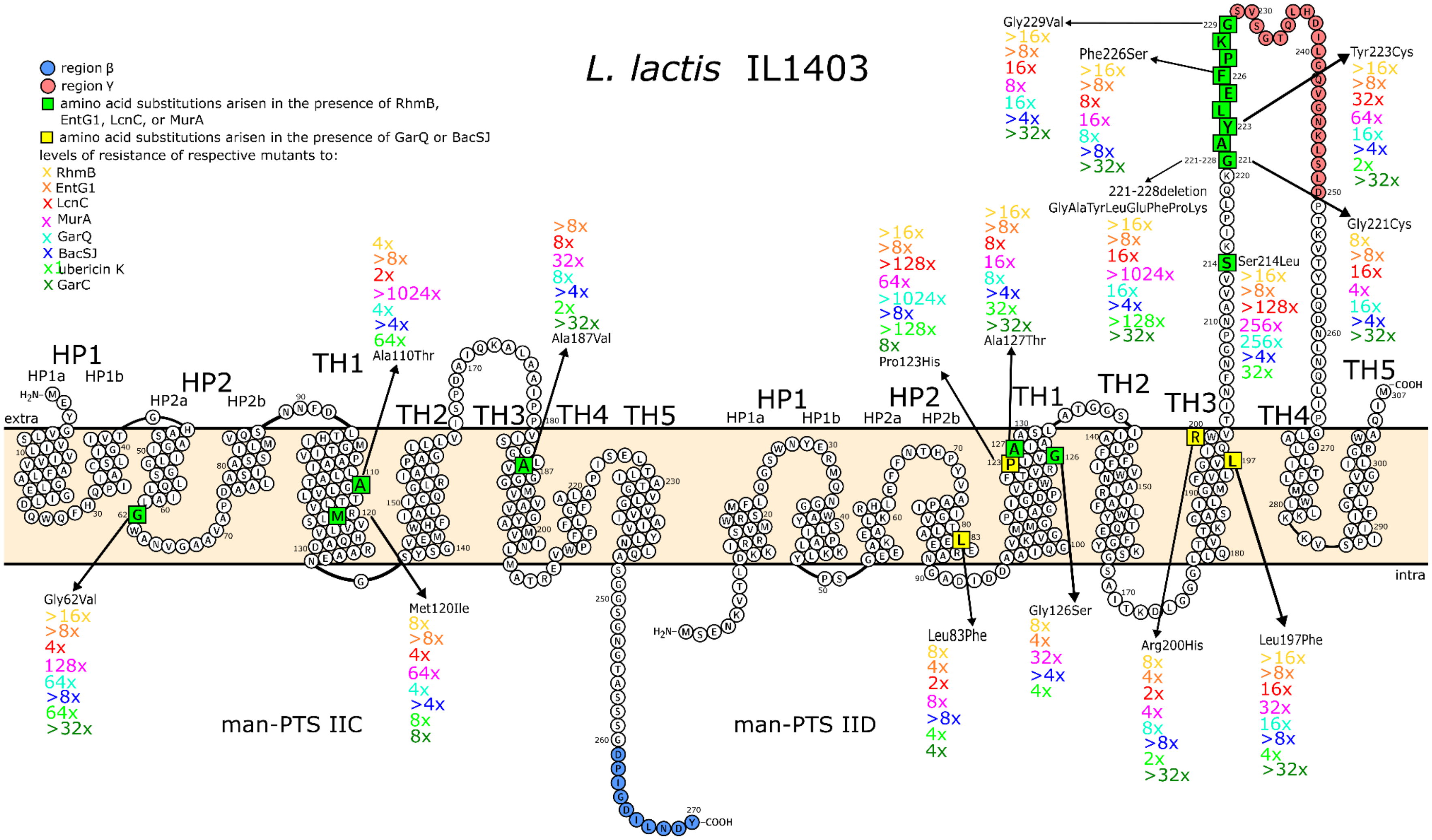

**(B)**

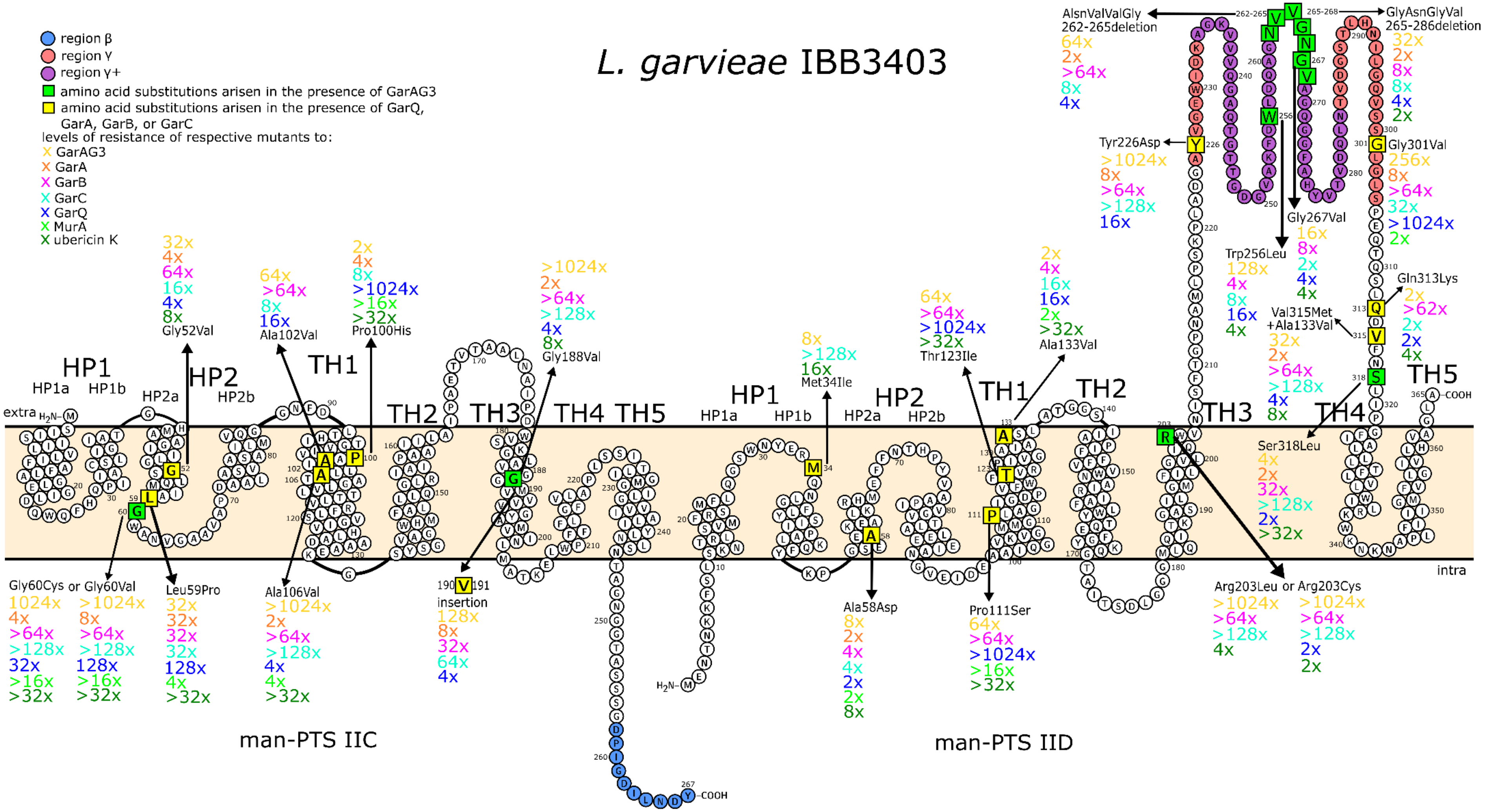
L. garvieae IBB3403
region β
region γ
region γ+
amino acid substitutions arisen in the presence of GarAG3
amino acid substitutions arisen in the presence of GarQ, GarA, GarB, or GarC
levels of resistance of respective mutants to:
X GarAG3
X GarA
X GarB
X GarC
X GarQ
X MurA
X ubericin K
HP1
HP1a
HP1b
HP2
HP2a
HP2b
TH1
TH2
TH3
TH4
TH5
extra
intra
Gly52Val
Ala102Val
Pro100His
Gly188Val
Met34Ile
Thr123Ile
Ala133Val
Gly60Cys or Gly60Val
Leu59Pro
Ala106Val
190 V 191 insertion
Ala58Asp
Pro111Ser
Arg203Leu or Arg203Cys
Ser318Leu
Val315Met +Ala133Val
Gln313Lys
Trp256Leu
Gly267Val
Gly301Val
Tyr226Asp
AlsnValValGly 262-265deletion
GlyAsnGlyVal 265-286deletion
man-PTS IIC
man-PTS IID

**Figure S7. Sequence alignment of GarQ-like peptides active against *L. garvieae* and *L. lactis* (A) or *L. lactis* only (B).** Fully conserved amino acids are in red, partially conserved ones—blue. Consensus symbols are: %—F or Y; !—I or V.

**(A)**

```
                     10        20        30        40        50        60        70
            ---------|---------|---------|---------|---------|---------|---------|-----
      GarQ                 EYHLMNGANGYLTRVNG--KYVYRVTKDPVSAVFGVISNGW-GSAGAGFGPQH
    GarAG2                 GTPLFYGANGYLTRENG--KYVYRVTKDPVSAVFGVISNGW-GSAGAGFGPQH
  LcnC/HB1  MVTTNKKSAPKKEPQFTKIMNSANGYLAYDNWNKKYVYHVTKDPVSAVAGVLANGW-GSAGAGFGPQTGGPSGKL
  MurA/HB7          MLKGKGYCKPVYYAANGYSCRYK-NGQWDYIVTKGNLEATLGVMSNGWVSSLGGGYFNRPK
Ubericin K             AKGVCKYVYPGSNGYACRYP-NGEWGYIVTKSNFEATKDVIVNGWVSSLGGGYFHGNRG
 Consensus  .....................gaNGYl.r.n.n.kyvY.VTKdpvsAv.gVisNGW.gSaGaG%gpq........
```

**(B)**

```
                     10        20        30        40        50        60        70
            ---------|---------|---------|---------|---------|---------|---------|--
    BacSJ                     YSYFGGSNGYSWRDKRGHWHYTVTKGGFETVIGIIGDGWGSAGAPGPGQH
 LcbC/HB14                     NFFGGSNGYSWRDKKGHWHYTVTSGVSSTVAQIIGNGWGSAGAPGVGQR
 RhmB/HB12                     VRIYGSNGYAWTDKHGHWHYTVTKGAVDTVFGIIGNGWGSAGASGRQH
  AglA/HB5  MAALLFFIKAAIRNSGYCEPVYYGANGYSCRYSNGKWDYKVTKTAAQAVGGVMAKGWIGSLGGSYYQPGYRG
   Angicin               GSGYCKPVMVGANGYACRYSNGRWDYKVTKGIFQATTDVIVKGW-AEYGPWIPRH
 EntG1/HB3          MRTIQGVG--RPIFNGANGYLSRDKYGHYTYTVTKGPLEAALGVMVNGRASTAGGGSWAPKYN
 Consensus  ...............g...p...GaNGY..rdk.Ghw.YtVTkg...av.g!i.nGw.s..g.g........
```

**Figure S8. CD spectra of (A) GarQ in water, 25% TFE or 50% TFE, and (B) its amino acid variants in 50% TFE**

**(A)**

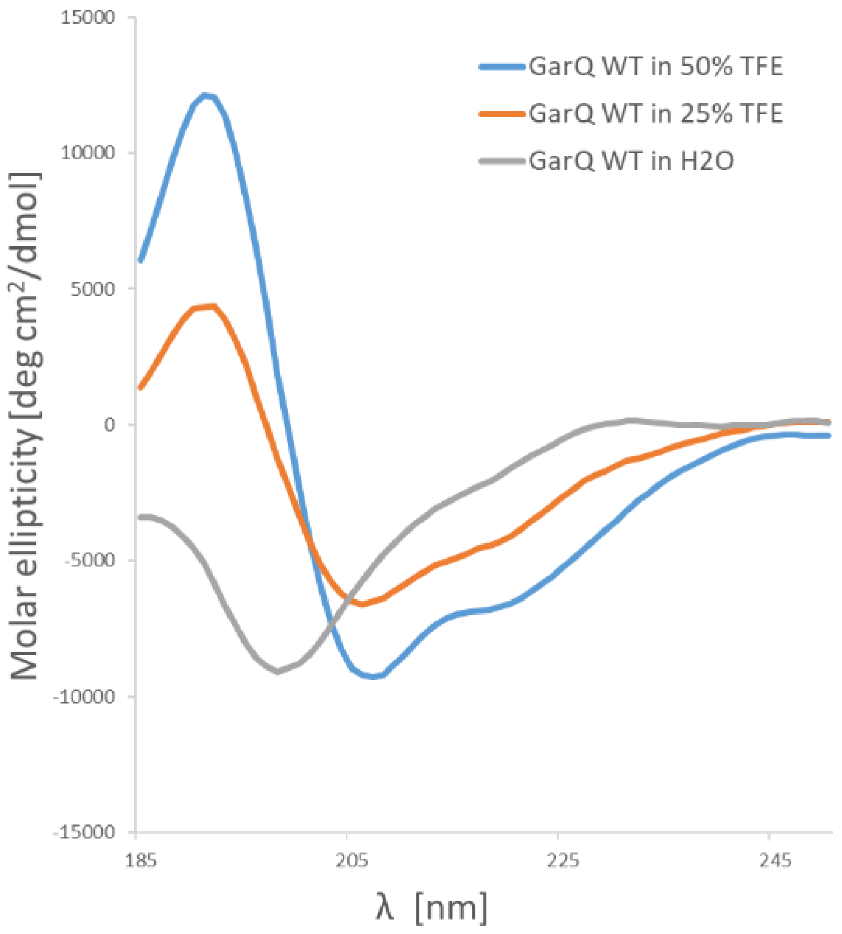


**(B)**

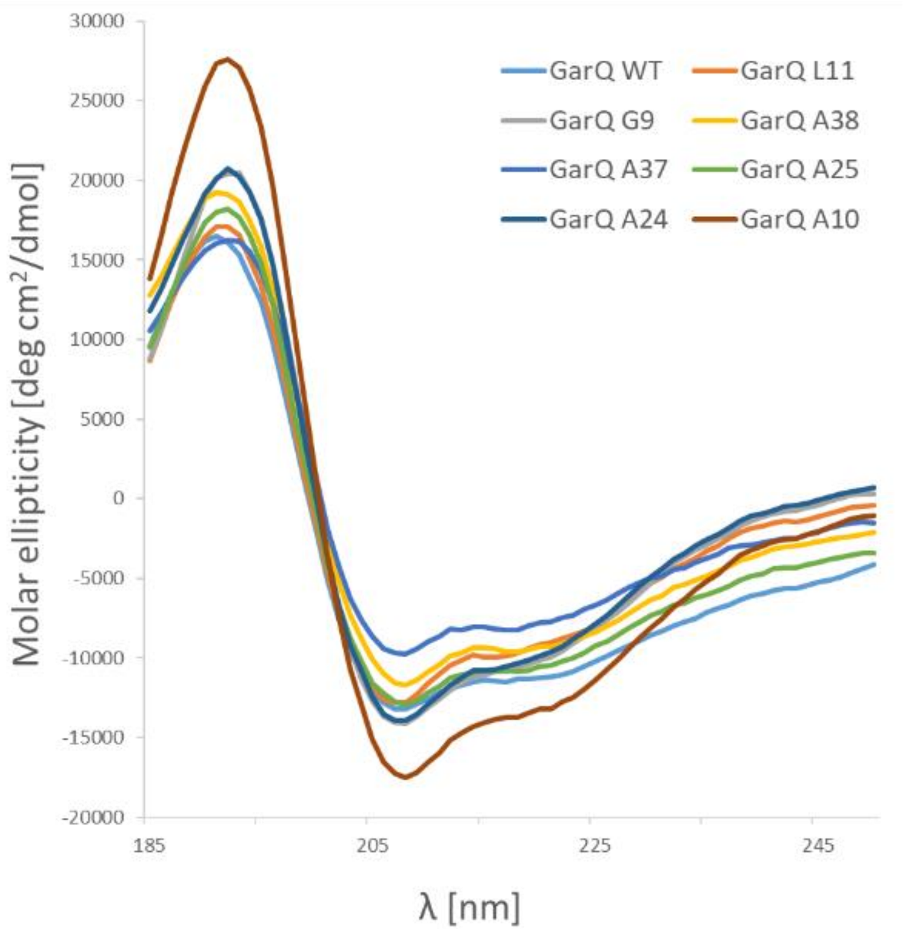

**Figure S9. Schematic presentation of the arrangement of secondary structures in the main representatives of Man-PTS-binding bacteriocins.** The proportions of the lengths of the peptides and their secondary structures have been preserved.

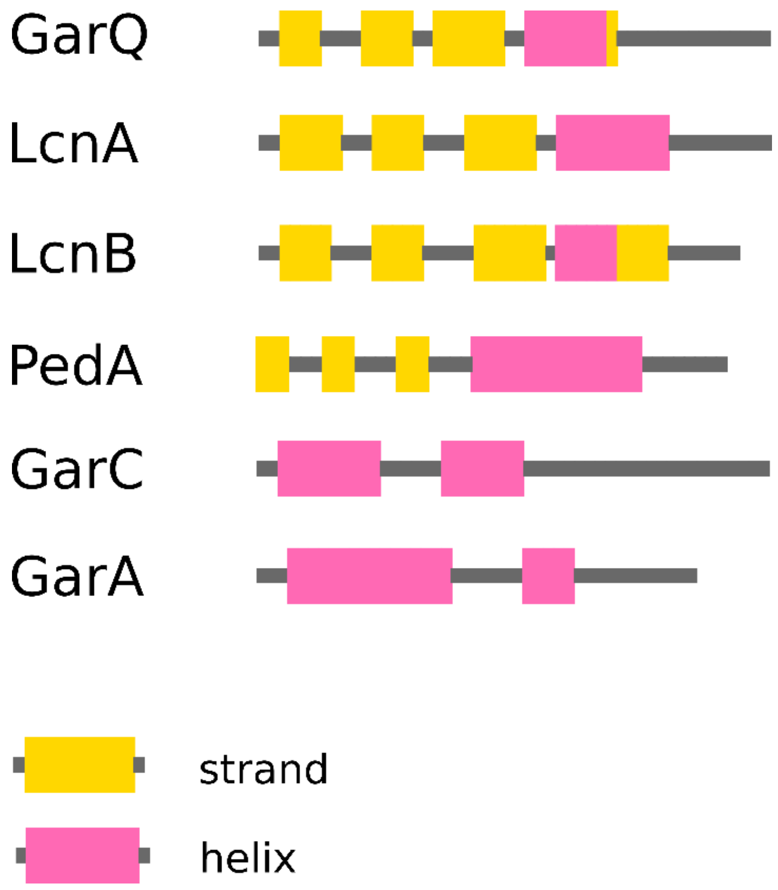
